# Converting non-neutralizing SARS-CoV-2 antibodies targeting conserved epitopes into broad-spectrum inhibitors through receptor blockade

**DOI:** 10.1101/2022.01.24.477625

**Authors:** Payton A.-B. Weidenbacher, Eric Waltari, Izumi de los Rios Kobara, Benjamin N. Bell, John E. Pak, Peter S. Kim

## Abstract

All but one of the authorized monoclonal antibody-based treatments for SARS-CoV-2 are largely ineffective against Omicron, highlighting the critical need for biologics capable of overcoming SARS-CoV-2 evolution. These mostly ineffective therapeutic antibodies target epitopes that are not highly conserved. Here we describe broad-spectrum SARS-CoV-2 inhibitors developed by tethering the SARS-CoV-2 receptor, angiotensin-converting enzyme 2 (ACE2), to antibodies that are known to be non-neutralizing, but which target highly conserved epitopes in the viral spike protein. These inhibitors, called <u>Re</u>ceptor-blocking <u>co</u>nserved <u>n</u>on-<u>n</u>eutralizing <u>A</u>nti<u>b</u>o<u>d</u>ies (ReconnAbs), potently neutralize all SARS-CoV-2 variants of concern (VOC), including Omicron. Neutralization potency is dependent on both the binding and inhibitory ReconnAb components as activity is lost when the linker joining the two is severed. In addition, a bifunctional ReconnAb, made by linking ACE2 to a bispecific antibody targeting two non-overlapping conserved epitopes, defined here, shows sub-nanomolar neutralizing activity against all VOCs, including Omicron. Given their conserved targets and modular nature, ReconnAbs have the potential to act as broad-spectrum therapeutics against SARS-CoV-2 and other emerging pandemic diseases.

## Introduction

The emergence of the Omicron variant has rendered six of the seven^1–9^ clinically available monoclonal antibodies (mAbs) essentially ineffective against SARS-CoV-2 – only sotrovimab retains robust neutralizing activity against Omicron^10, 11^. These clinical mAbs all target the receptor binding-domain (RBD)^1–9^ of the spike protein and were selected for their neutralizing potency against Wuhan-Hu-1 SARS-CoV-2. The six mAbs besides sotrovimab target non-conserved (variable) regions of the RBD^4, 12–18^ and prevent interaction with its receptor, human angiotensin converting enzyme 2 (ACE2)^16, 19–21^. Sotrovimab, a derivative of the mAb S309^22–25^, was initially isolated from a SARS-CoV-1 survivor, so its epitope in the RBD is more highly conserved,^9^ although *in vitro* escape mutations have been identified^14^.

The spike protein is large (∼450 kDa as a trimer), and contains extensive regions that are extremely highly conserved (Fig. 1A)^26–29^. Some residues on the spike that are distant from the RBD have near-perfect sequence identity within related coronaviruses (Fig. 1B)^30, 31^. Presumably, these regions are highly conserved because they are required for viral activity (e.g., membrane fusion)^32^. While the spike protein of Omicron has a much larger mutational profile than that of previous variants of concern (VOCs)^33^, with 36 total mutations, 15 being in the RBD^11, 34, 35^, the highly-conserved epitopes remain largely unaltered (Fig. 1).^11, 34, 35^

**Figure 1.**
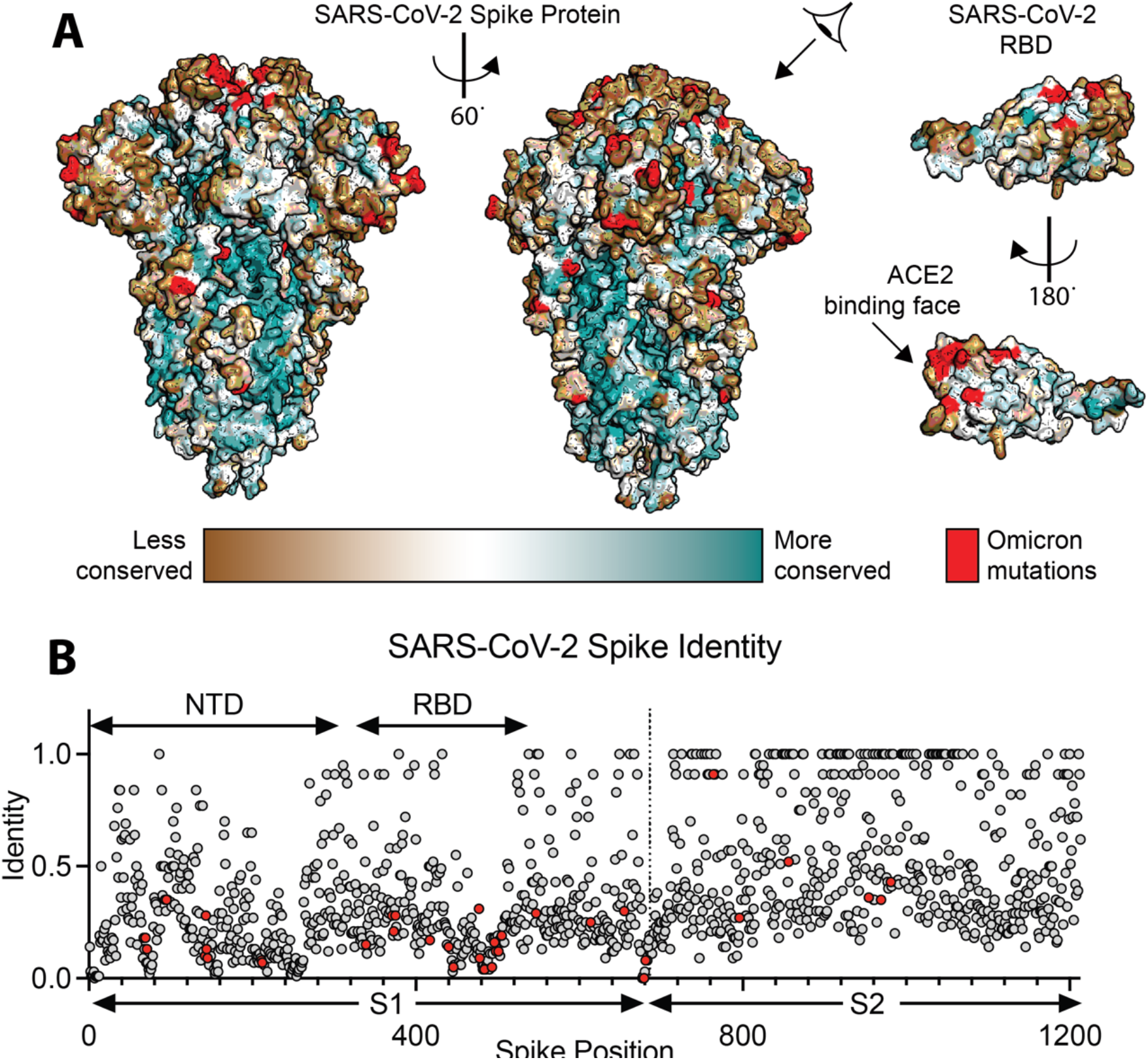
Conservation of the SARS-CoV-2 spike protein. (A) Sequence conservation from 44 related spike proteins overlaid on the SARS-CoV-2 spike protein structure (left) and the SARS-CoV-2 RBD (right – residues 319-541) (PDB ID: 6VXX) identifies a highly conserved patch in S2. (B) Sequence identity for all residues in the SARS-CoV-2 spike protein compared to a set of 44 related coronavirus spike proteins shows higher conservation in the S2 relative to S1. A value of 1.0 means perfect identity across all compared coronavirus proteins. RBD and NTD domains of SARS-CoV-2 spike are labeled on top, S1 and S2 domains are labeled on bottom.

In other viral spike proteins, for instance influenza hemagglutinin^36–41^, highly conserved epitopes outside of the receptor-binding region are targets of potent broadly neutralizing antibodies (bnAbs). However, despite the heightened interest sparked by the global pandemic, the search for bnAbs against betacoronaviruses has been largely disappointing. While one conserved helical epitope at the base of the spike protein has been shown to elicit rare mAbs with relatively broad neutralizing activity, their potency is often weaker than RBD-directed neutralizing Abs^42–45^. Further, neutralizing N-terminal domain (NTD)^46, 47^ antibodies have been identified, but their epitopes are not highly conserved.

Indeed, available evidence suggests that conserved regions outside the RBD generally elicit non-neutralizing mAbs^43, 48–54^. We hypothesized that we could generate potent, broad-spectrum inhibitors by modifying existing non-neutralizing antibodies which target highly conserved epitopes on the spike protein to also contain a receptor-blocking component. Due to the conservation of their epitopes, such inhibitors would potentially be broadly neutralizing.

Here we introduce <u>Re</u>ceptor-blocking <u>co</u>nserved <u>n</u>on-<u>n</u>eutralizing <u>A</u>nti<u>b</u>odie<u>s</u>, (ReconnAbs pronounced recon-abs), a novel class of therapeutic proteins in which non-neutralizing antibodies that target highly conserved, non-RBD epitopes, are tethered to the ACE2 receptor, which otherwise has low intrinsic affinity and neutralizing potency. The cross-reactive, non-neutralizing antibodies were identified in a two-step process. First, we analyzed the phylogenetic trees of a collection of SARS-CoV-2 antibodies and eliminated those that are likely to bind the RBD. Then, similar to the development of sotrovimab,^9^ we determined which of these non-RBD antibodies bound to the SARS-CoV-1 spike protein. We predict that ReconnAbs will have both increased potency, due to the increase in effective concentration of each component^55^, as has been shown previously for non-neutralizing antibody fusions^56^, but more importantly, increased broad-spectrum activity by targeting highly conserved, non-RBD epitopes on spike. ReconnAbs show neutralizing activity against all SARS-CoV-2 VOCs tested, including Omicron. Further, a bispecific ReconnAb containing two non-neutralizing antibodies with non-overlapping epitopes fused to ACE2 confers sub-nanomolar neutralization against all VOCs tested. Our findings reveal the benefit of repurposing highly cross-reactive, non-neutralizing antibodies to create a new class of broad-spectrum anti-viral agents.

## Results

To first profile the landscape of non-neutralizing antibodies, we produced a library of SARS-CoV-2 spike-binding antibodies not directed against the RBD. To ensure library diversity, we first curated the publicly available SARS-CoV-2 antibody repository CoV-AbDab^46, 57–60^ for sequences specifically from COVID-19 convalescent donors that bound to the spike protein outside of the RBD. From this limited set of 696 antibody sequences, we generated phylogenetic trees for both the antibody heavy chain (HC) and light chain (LC). We constructed phylogenies using amino acid sequences of full-length V-genes and the CDR3 region. We also included one allele of each germline V-gene into the phylogenies to provide context for germline diversity of the antibody dataset.

We compiled non-RBD-binding antibodies, focusing specifically on the clustering of the non-RBD-binding sequences within the HC phylogenetic tree (Fig. 2A). We identified distinct clusters on the heavy/light chain phylogenetic trees and chose 48 diverse sequences spread throughout the tree (Fig. 2A and Supplementary Fig. 1). These included at least one antibody sequence from all clusters containing at least four or more non-RBD-binding antibodies. The sequences we chosen based on their HC sequences also showed diversity in the LC phylogenetic tree (Supplementary Fig. 1). These 48 clones display a range of CDRH3 and CDRL3 lengths (Fig. 2B) and utilize an array of V genes in both the HC and LC (Fig. 2C), further confirming their diversity.

**Figure 2.**
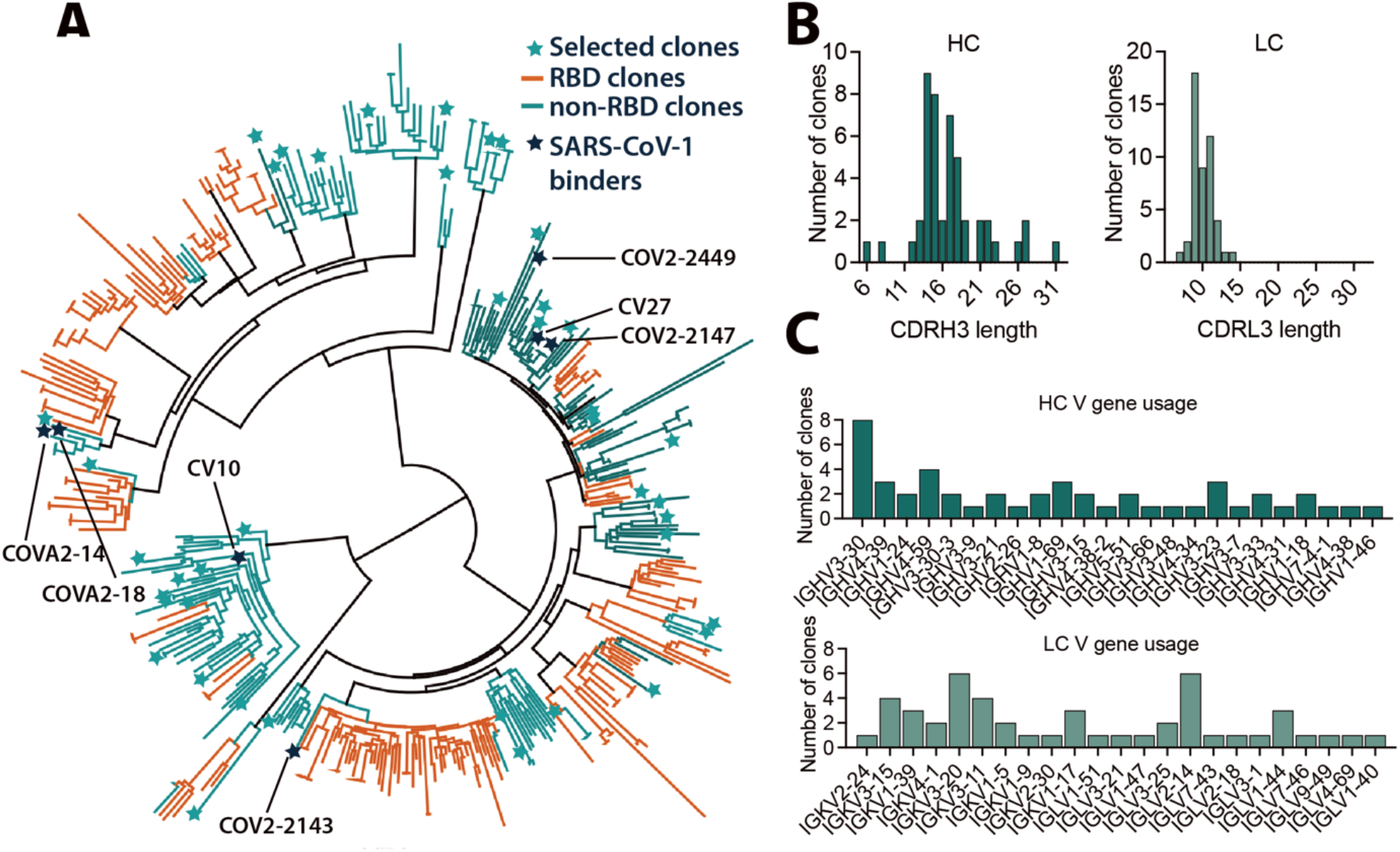
non-RBD antibodies were selected to prioritize diversity. (A) A phylogenetic tree of 422 heavy chain sequences, from our curated library of 696 anti-SARS-COV-2 spike antibodies, was generated using Geneious Prime. Germline alleles not shown. 48 selected clones shown as stars. Histograms of the HC and LC (B) CDR3 lengths (C) V gene usage from the 48 selected non-RBD clones indicated in (A).

To determine which of these 48 non-RBD-binding antibodies target highly conserved epitopes, we used binding to the SARS-CoV-1 spike as a surrogate for epitope conservation. We designed the 48 scFvs constructs by fusing the antibody HC and LC variable regions to the yeast surface protein Aga2p^61–63^ to enable yeast-surface display. To profile the scFv panel, we optimized production of biotinylated SARS-CoV-2 and other human coronavirus (hCoV) spike proteins (Supplementary Fig. 2A-D) and produced biotinylated versions of the SARS-CoV-2 and SARS-CoV-1 spike proteins. These were used to probe the yeast library by fluorescence flow cytometry (Fig. 3A). The complete 48-member library showed robust (82%; Fig 3A) staining with the SARS-CoV-2 spike, consistent with the original antibody collection having been isolated from SARS-CoV-2 convalescent donors. The library had reduced (21%; Fig 3A) staining with the SARS-CoV-1 spike. Consistent with the intention of the library, no clones bind to the RBD of SARS-CoV-2 (Fig. 3A).

**Figure 3.**
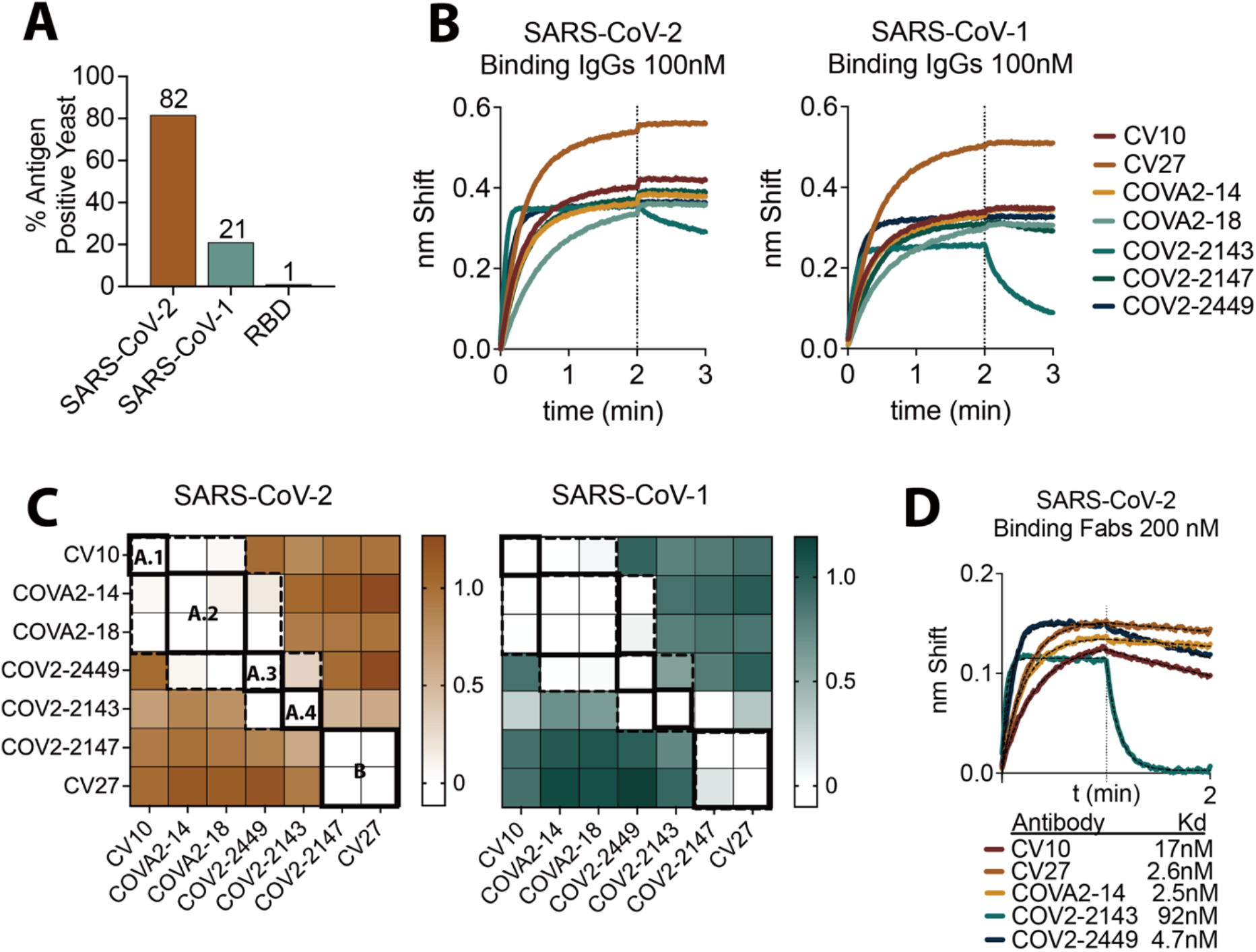
A subset of non-RBD SARS-CoV-2 antibodies bind SARS-CoV-1, a surrogate for epitope conservation. (A) A binding profile of the scFv-yeast library produced from the sequences identified in Fig. 1 and Supplementary Fig. 1. (B) BLI binding of identified cross-reactive clones expressed as IgGs binding at 100 nM to SARS-CoV-2 spike (left) or SARS-CoV-1 spike (right). (C) BLI competition binding assay of the seven cross-reactive antibodies binding to SARSCoV- 2 (left) and SARS-CoV-1 (right). White indicates no binding of the tested antibody, demonstrating the antibodies compete for binding. Antibodies which compete are surrounded by dotted lines, unique competition groups are surrounded by solid lines. The 5 unique competition groups are labeled on the SARS-CoV-2 binding competition map. Site A.1-A.4 is indicated as an overlapping supersite. Loading antibodies are indicated in columns and competing antibodies are indicated in rows. (D) Binding of antibodies FAb fragments at 200nM against SARS-CoV-2 spike. Hashed lines show KD fit determined using Prism.

Having identified 48 antibodies that bind outside the RBD, we next selected those that bind to highly conserved regions of the spike protein. To do this, we used fluorescence-activated cell sorting (FACS)^64^ with the SARS-CoV-1 spike protein as bait and identified ten sequences. We confirmed that the corresponding full-length IgG antibodies (Supplementary Fig. 3) bind to both SARS-CoV-2 and SARS-CoV-1 spike proteins by ELISA (Supplementary Fig. 4). Of the ten, seven clones were strong SARS-CoV-1 binders, confirmed by biolayer interferometry (BLI) (Fig. 3B). One clone, COV2-2449, also binds MERS and OC43 spike proteins (Supplementary Fig. 4 and 5). Consistent with previous reports^4, 57–60^, these antibodies were non-neutralizing in our assay (Supplementary Fig. 6).

**Figure 5.**
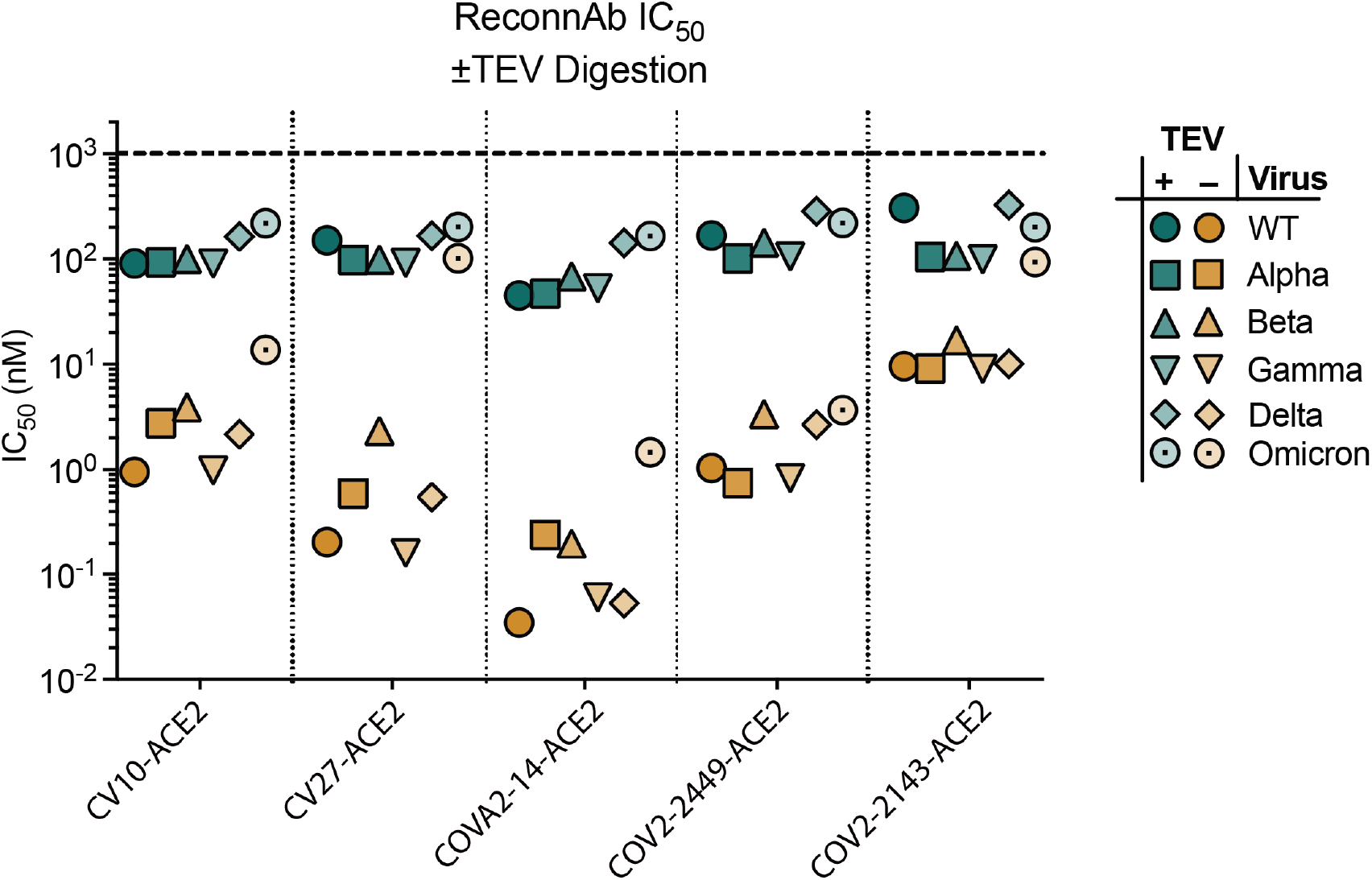
Intact ReconnAbs (orange) show broad spectrum neutralization of SARS-CoV-2 VOCs. Inhibition is markedly reduced upon TEV cleavage (teal). Pseudoviral 50% inhibitory concentration (IC50) for ReconnAbs (bottom) against a range of SARS-CoV-2 VOCs with and without TEV cleavage. IC50 values shown are the average of two independent experiments.

We used BLI to characterize the epitopes of these seven antibodies in a binding competition assay. We loaded each antibody onto either SARS-CoV-2 or SARS-CoV-1 spike proteins, and then we tested for subsequent binding of each of the other antibodies (Fig. 3C). The results suggest that there are five primary epitopes, of which four are in a partially overlapping supersite (Fig. 3C) – likely corresponding to the extensive, continuous patch of highly conserved residues on the spike protein surface (Fig. 1A). Two sets of antibodies had identical epitopes – the pair^60^ of COVA2-14 and COVA2-18, and the pair of CV27 and COV2-2147. This result is consistent with the phylogeny, which shows the antibodies in the two pairs clustered very closely together (Fig. 2A and Supplementary Fig. 1). The identification of five unique epitopes from the seven selected antibodies highlights the diversity in the initial starting library.

We selected five antibodies, one from each described epitope (Fig. 3D), and converted these non-neutralizing, cross-reactive antibodies into ReconnAbs by fusion to the ACE2 ectodomain, as the receptor-blocking component of the ReconnAb design. We designed the linker to be long enough to allow for simultaneous binding of both ACE2 to the RBD and the scFv, regardless of epitope, to the spike S2 domain (Fig. 4A and Supplementary Fig. 7). We joined the C-terminus of the scFv to the N-terminus of ACE2, because the N-terminal residue of the ACE2 ectodomain is adjacent to the SARS-CoV-2 RBD when bound. We also incorporated within the linker a hexa-histidine tag for purification and a TEV protease site to enable assessment of ReconnAb activity when its binding and inhibitory components are separated (Fig. 4A and Supplementary Fig. 7). We anticipated that ReconnAbs would bind to both a highly conserved site on the spike protein and simultaneously to the RBD through the ACE2 domain (Fig 4B). However, if cleaved at the TEV site, the intrinsically low-affinity ACE2 domain would not benefit from the affinity of the non-neutralizing antibody (Fig. 4B).

**Figure 4.**
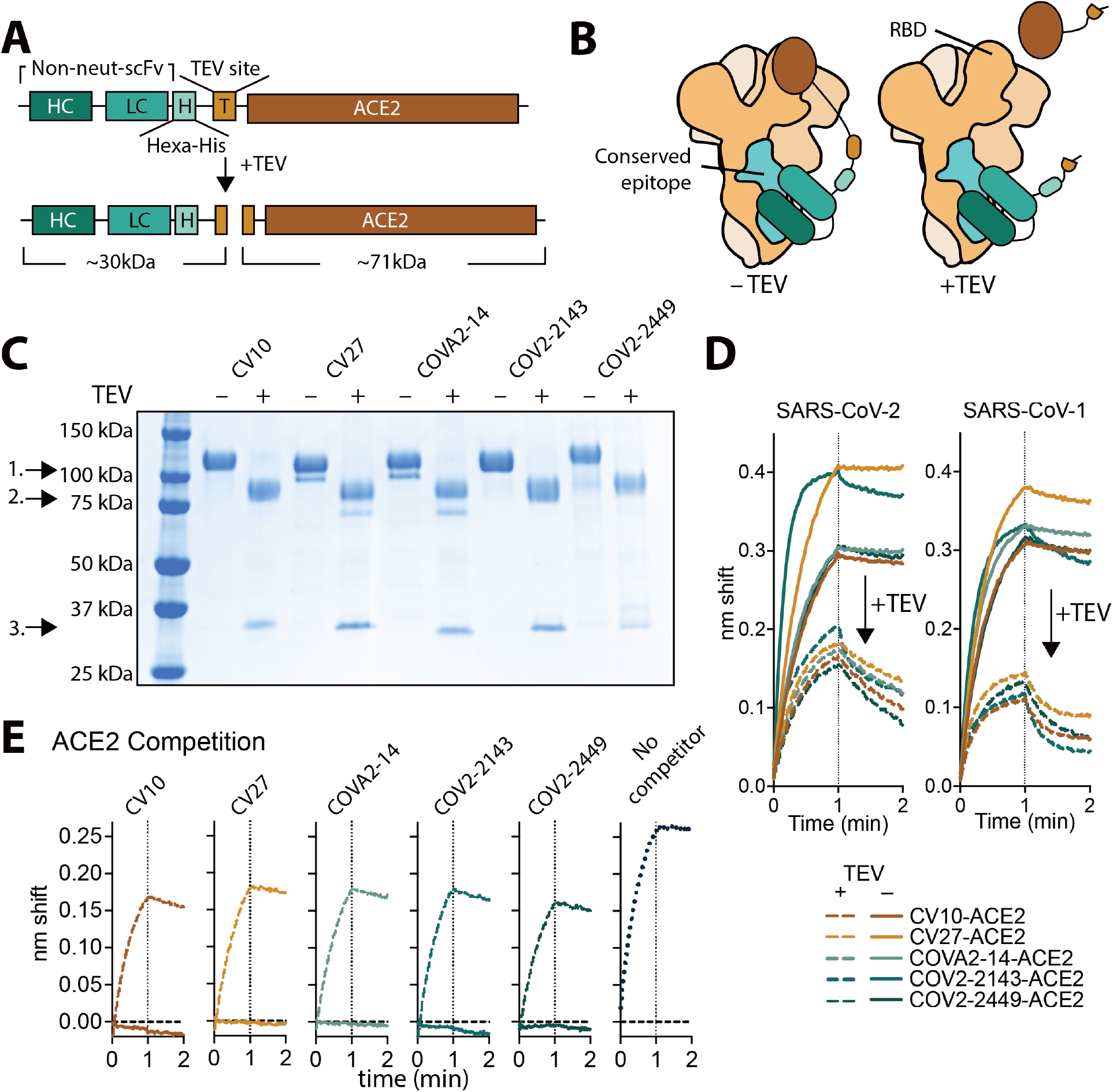
scFv-based ReconnAbs tether ACE2 to cross-reactive, non-neutralizing antibodies. (A) A schematic of an scFv-based ReconnAb protein before and after TEV cleavage. Estimated molecular weights of cleavage products are shown beneath both. (B) schematic of ReconnAb activity and the dependence on the tether. The scFV binds to a conserved site and then ACE2 interacts with the RBD. Upon TEV cleavage, the ACE2 has lower apparent affinity and does not bind the RBD (right). Conserved sites shown as teal, remainder of spike monomers shown as tints of brown. (C) SDS-PAGE demonstrates ReconnAbs are readily cleaved by TEV. 1. Full-length ReconnAb, 2. ACE2, 3. scFv. (D) BLI binding of uncleaved and TEV-cleaved ReconnAbs to either SARS-CoV-2 spike (left) or SARS-CoV-1 spike (right) show a reduction in binding upon TEV cleavage. (E) BLI binding of hFc-ACE2 to SARS-CoV-2 spike which have been pre-associated with ReconnAbs either uncleaved (solid lines) or cleaved (hashed lines) show TEV-cleaved ReconnAbs do not compete with hFc-ACE2 binding. Binding of hFc-ACE2 without competitor shown on the right (dotted line).

We expressed and purified the five ReconnAbs and used gel electrophoresis to confirm that TEV cleavage separated the ACE2 and scFv components (Fig. 4C). BLI experiments showed that TEV cleavage of the ReconnAb proteins reduced binding to both SARS-CoV-2 and SARS-CoV-1 spike proteins (Fig. 4D), consistent with the lower affinity of monomeric ACE2^65^. We then investigated the ability of the ReconnAbs to block ACE2 binding to the SARS-CoV-2 spike protein. ACE2 competition is often used as a surrogate for neutralization, as preventing ACE2 binding prevents the virus from interacting with target cells^66^. Indeed, uncleaved ReconnAbs show substantial interference with binding of an Fc version of human ACE2 (hFc-ACE2) whereas TEV- cleaved ReconnAbs do not (Fig. 4E).

We next investigated if ReconnAbs were able to neutralize lentiviral pseudoviruses corresponding to the SARS-CoV-2 VOCs and found that all ReconnAbs neutralized all VOCs – some showing nanomolar potency against Omicron (Fig. 5). Consistent with its lower affinity (Fig. 3D), COV2-2143-ACE2 had the weakest neutralization of the tested ReconnAbs (Fig. 5). COV2- 2449-ACE2 showed the least deviation in neutralization potency between variants, consistent with its epitope being the most highly conserved (Supplementary Fig. 4 and 5). Importantly, the TEV- proteolyzed versions of the ReconnAbs did not confer the same neutralizing potency as their uncleaved counterparts (Fig. 5), demonstrating that the separate components are not working synergistically, but that the tether is essential for the ReconnAb components to work cooperatively.

Two ReconnAbs, CV10-ACE2 and COV2-2449-ACE2, were of particular interest as they showed broad-spectrum neutralization (Fig. 5) and did not have overlapping epitopes (Fig. 3C). We postulated that a bifunctional IgG-ReconnAb containing both CV10 and COV2-2449 would make viral escape even less likely. To produce a bifunctional IgG-ReconnAb, we utilized the CrossMAb platform^67–69^ and tethered ACE2 to the LC of only one of the IgG arms (Fig. 6A, B). This allows, as with the scFv-ACE2 fusions, that the stoichiometry be only a single ACE2 per ReconnAb, such that ACE2 remains monovalent before and after TEV-cleavage.

**Figure 6.**
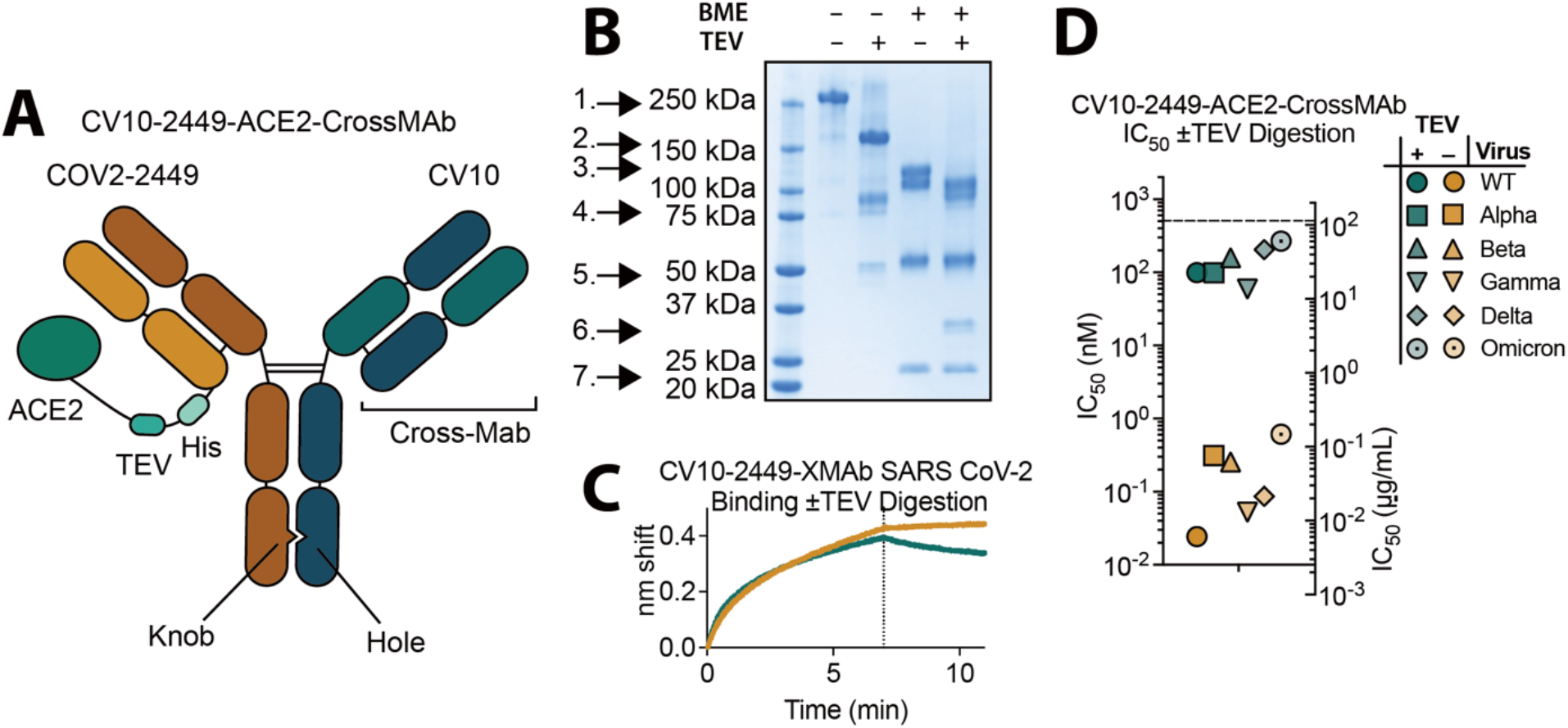
A bifunctional IgG ReconnAb shows potent neutralization of SARS-CoV-2 VOCs. (A) A schematic representation of the CV10-2449-ACE2-CrossMAb indicates linkage of ACE2 and bifunctional Fab arms. (B) The CV10-2449-ACE2-CrossMAb ReconnAb contains all designed components by SDS-PAGE, assayed before or after TEV cleavage and with or without 2-mercaptoethanol (BME). 1. Full-length ReconnAb, 2. Cleaved CrossMAb CV10-2449 IgG, 3. COV2-2449-LC-ACE2 fusion, 4. ACE2 (reduced ACE2 shows double banding), 5. HC, 6. Cleaved 2449-LC with linker, 7. CV10 LC. (C) BLI binding of CV10-2449-ACE2-CrossMAb (brown) or the TEV cleaved form (teal) to SARS-CoV-2 spike. Binding is reduced upon TEV cleavage. (D) Pseudoviral 50% inhibitory concentration (IC50) for CV10-2449-ACE2-CrossMAb against a range of SARS-CoV-2 VOCs with and without TEV cleavage. IC50 values shown are the average of two independent experiments.

We expressed and purified the CV10-2449-ACE2-CrossMAb (Fig. 6B) and found that it bound to SARS-CoV-2 as expected (Fig 6C). As well, the uncleaved CrossMAb competed substantially with ACE2 (Supplementary Fig. 8). Dependent on the presence of the tether, the CV10-2449-ACE2-CrossMAb neutralized all SARS-CoV-2 VOCs, including Omicron, at sub-nanomolar concentrations (Fig. 6D). Taken together, the results described here lay a foundation for the development ReconnAbs as a novel class of broadly neutralizing therapeutics.

## Discussion

Traditionally, the discovery of therapeutic biologics against infectious diseases has focused on identifying agents with neutralizing activity. We demonstrate here using ReconnAbs, that cross-reactive, non-neutralizing antibodies – which have been often largely overlooked – can be powerful reagents in the creation of potent, broad-spectrum anti-viral agents.

ReconnAbs have two main components: a binding component – the non-neutralizing antibody that binds with high-affinity to a conserved region on the spike protein – and an inhibitory component, in our case, the ACE2 domain^70, 71^. Since therapeutics containing ACE2 run the risk of eliciting autoimmunity in humans, our use of ACE2 as the inhibitory component represents a proof-of-concept of the ReconnAb design. The ACE2 module could be replaced by other neutralizing components such as ACE2 domains with enhanced RBD-binding activity^72–74^, aptamers^75, 76^, or RBD-directed mAbs^1–9, 46, 56–60, 77^. Future ReconnAb designs could also target the interaction with dipeptidyl peptidase 4 (DPP4), a receptor for other coronaviruses^78^, furthering their breadth.

Improvements could also be made to the conserved, non-neutralizing antibodies. Our library of SARS-CoV-2 non-RBD antibodies was derived from sequences early in the COVID-19 pandemic, which is relatively small in scope. The library does not contain, for instance, any vaccine-derived antibodies, which are known to include cross-reactive, non-neutralizing antibodies^49^. Future iterations of this work could start with much larger libraries^57^, with the potential to identify antibodies and/or nanobodies that target the most highly conserved epitopes and that are least likely to undergo mutational escape^79^. Although we have focused on non-neutralizing antibodies, neutralizing antibodies that bind to conserved epitopes might also be useful as the conserved component of ReconnAbs. Other features like linkage locations and length, fusion partners, and modifications to the Fc domains can be tuned in subsequent ReconnAb designs, and will likely play an important role in their future conversion to therapeutics^23–25^.

Compared to neutralizing epitopes, highly conserved, non-neutralizing epitopes are less likely in theory to be subject to immune pressure since their mutations do not affect the ability of the virus to infect cells. Omicron provides strong evidence that SARS-CoV-2 viral evolution responds to immune pressure by mutating neutralizing epitopes (Fig.1)^10, 11, 34, 35^. ReconnAbs demonstrate the powerful utility of a non-active component, if it targets a highly conserved epitope, in the development of therapeutics.

Finally, we anticipate that interrogation of existing antibody libraries for highly conserved, non-neutralizing binders will facilitate production of ReconnAbs, not just for SARS-CoV-2 but also for other viruses like HIV-1, influenza^80^, or other human coronaviruses. We see ReconnAbs as having utility not only in the current pandemic, but also in mitigating the impact of future pandemics. Strategic stockpiles of customized ReconnAbs and rapid administration in a pandemic setting could alleviate the initial impact of a new pathogen, allowing time for other therapeutics and countermeasures to be put into place.

## Acknowledgments

We thank J. Bloom and A. Greaney for plasmids and cells related to viral neutralization assays, J. DeRisi, K. Zorn, L. Matthew and M. Ott for Omicron-related plasmids, and I. Anderson at Stanford PAN facilities for rapid production of oligos utilized throughout this work. We are also grateful to T. Bruun and M. Filsinger Interrante and other members of the Kim Lab for fruitful discussions and helpful comments on earlier versions of this manuscript. The CMV/R expression vectors for IgG production were received from the NIH AIDS Reagent Program. This work was supported by the Virginia & D.K. Ludwig Fund for Cancer Research, the Frank Quattrone and Denise Foderaro Family Research Fund, and the Chan Zuckerberg Biohub.

## Methods

### Determination of sequence conservation in SARS-CoV-2 spike

Sequences of 42 of spike proteins, with between 30%-90% sequence conservation compared to SARS-CoV-2, as well as RaTG13 were aligned using the 6VXX sequence^26^ as a template. The sequences were aligned using Clustal Omega^30^ to develop a multiple sequence alignment (MSA). The MSA was uploaded onto the Consurf server^27–29^ for overlay onto the 6VXX structure on chain A. The resultant chain A was recolored based on conservation and replicated to replace chains B and C. The sequence alignment was again produced using MUSCLE^31^ and sequence identity was calculated using Geneious (Geneious Prime 2022.0.1). The alignment was truncated at residue 1213 where the sequence alignment dropped to only 9 sequences. The sequences used were: 6VXX, UniRef90_U5NJG5, _L7UP8, _A0A7U3W1C7, _K9N5Q8, _A0A2I6PIW5, _A0A3Q8AKM0, _U5WHZ7, _A0A5H2WTJ3, _A0A0U1WJY8, _A0A166ZND9, _A0A678TRJ7, _A0A2R4KP93, _A0A2Z4EVK1, _A0A7R6WCE7, _E0ZN36, _A0A6M3G9R1, _F1DAZ9, _A0A0U1UYX4, _A0A2R3SUW7, _A0A2Z4EVN5, _A0A2Z4EVN2, _U5LMM7, _A0A5Q0TVR4, _E0XIZ3, _A0A023Y9K3, _A0A2R4KP86, _A0A088DJY6, _A0A7G6UAJ9, _S4X276, _A0A4Y6GL90, _A3EXG6, _F1BYL9, _E0ZN60, _A0A0K1Z054, _A0A0U1WHI2, and NCBI accession numbers: YP_009047204.1, QLR06867.1, AAK32191.1, AGZ48828.1, AAT84362.1, QHR63300.2, ABD75513.1, YP_003767.1.

### Library Design

A library of antibodies directed against SARS-CoV-2 Spike (S) protein was developed using paired antibody sequences, meaning antibody sequences for which the heavy and light chain are both known, from the Coronavirus Antibody Database, CoV-AbDab^57^. All antibody sequences from convalescent COVID-19 donors which had been deposited before July 9^th^ 2021, were inserted into a table, categorized by their binding to the SARS-CoV-2 RBD portion of the spike protein or to a non-RBD portion of SARS-CoV-2 spike. Antibodies which were cataloged for non-RBD binding were preferentially identified, resulting in a total of 385 paired antibody sequences. For these non-RBD binding antibodies, the amino acid sequences of the corresponding heavy chain and light chain V-genes and CDR3 regions, already compiled from the Coronavirus Antibody Database, were imported into Geneious Prime v2021.1.1 (a bioinformatics software; geneious.com). Using Geneious Prime, the heavy chain sequences and light chain sequences were separately analyzed to produce phylogenetic trees. For these phylogenetic trees, RBD binding antibodies were also included to ensure selection of antibody sequences that were both non-RBD binding and clearly distinct from RBD-binding sequences. 371 RBD-binding antibody, 59 germline antibody, and 325 non-RBD binding antibody nucleic acid sequences of the corresponding heavy chain and light chain genes were imported, for a total of 755 heavy chain and light chain sequences (696 excluding germline antibodies). The sequences were first aligned using the MUSCLE algorithm, and then two phylogenetic trees were made, both using PhyML 3.3.20180621. The sequence similarities used to produce phylogenetic trees account for antibody germlines, CDR lengths, and amount of somatic hypermutation. After producing phylogenetic trees based on the heavy chain and light chain sequences, a total of 48 sequences were identified based on their location in the phylogeny. Distinct clusters, composed of only non-RBD sequences, on the heavy chain phylogenetic trees were noted, and a single representative sequence was selected from each, chosen to also include distinct light chain sequences whenever possible.

### scFv design

The sequences of these 48 antibodies were then converted into scFv sequences by linking the HC variable region to the LC variable region with a G4S-3 linker (GGGGSGGGGSGGGGS). All scFvs were designed in the order: signal sequence-HC-G4-S-3-LC. This vector also contained the HVM06_Mouse Ig heavy chain V region 102 signal peptide (MGWSCIILFLVATATGVHS) to allow for protein secretion and purification from the supernatant. Following construct design, the plasmids were ordered with the sequences inserted at the XhoI and NheI sites in the pTwist CMV BetaGlobin vector (Twist Biosciences).

### Library Production

4µg of pPNL6 vector in Cut Smart buffer was digested using 1µl of NheI HF and BamHI HF (NEB Biolabs) at 37°C for 1h. Digested plasmid was then gel extracted using Thermofisher Scientific Gel Extraction Kit. Equimolar aliquots of each scFv plasmid were pooled and the resultant pool was amplified using primers which annealed to the hexa-his Tag (reverse primer) or signal peptide (forward primer) and had a 50bp overlap with the pPNL6 vector digested with NheI and BamHI. The pooled amplification was gel extracted to ensure it was the correct size. Yeast were prepared by first streaking a YPAD plate and incubating for 2-3 days until a single colonies were identifiable. A single colony was inoculated in 5 mL of YPAD overnight shaking at 30°C. Cultures were harvested into 6 tubes and pelleted. Yeast were resuspended in electroporation buffer (10 mM Tris Base, 250 mM sucrose, 2mM MgCl2) containing the gel extracted library amplification and digested pPNL6 vector. This mixture was then pulsed and the electroporated yeast were recovered in SD-CAA media overnight (30°C shaking). These yeast were then induced by a 1:10 dilution into SG-CAA media and grown at 20°C shaking for 2-3 days.

### Yeast Binding

Following induction in SG-CAA shaking for 2-3 days at 20°C, the yeast library, expressing surface exposed scFvs, was incubated for 15 mins with a dilution of preformed baits. Baits were formed by mixing biotinylated baits and streptavidin 647 (Jackson Immunoresearch) at a 4:1 ratio. For example, 250nM bait would be produced by incubation of 250 nM biotinylated antigens and 62.5 nM streptavidin 647. Yeast were flowed with two colors of “bait,” the first (FITC) stains for a c-myc tag. The c-myc tag is a surrogate for expression as the scFv constructs in the pPNL6 vector contain an in-frame C-terminal c-myc tag. So, any yeast which are c-myc positive are displaying full-length antibodies. The second color bait (Alexa Flour 647 – APC channel) stains for the antigen-target of the scFv. To make the stain, streptavidin with an Alexa Flour 647 tag is incubated with biotinylated bait protein. This complex is then used to stain the yeast. Any yeast which are positive for Alexa Flour 647, are then binding to the protein antigen. Yeast were spun down and resuspended in in 50 µl PBSM containing the respective concentration of tetrameric bait. After 15mins cells were then washed 1x with PBSM and then resuspended in 50µL PBSM containing 1 µl of anti-c-myc FITC (Miltenyi) for 15 mins. Samples were then washed 2x with PBSM and then resuspended in 50 uL of PBSM. These samples were flowed (Accuri C6 flow cytometer) and the percent antigen positive was determined as the ratio of antigen positive cells divided by all cells expressing scFv (c-myc positive) multiplied by 100. Gates were set such that ∼.5% of yeast were antigen positive in the streptavidin alone control.

### Yeast Sorts

The yeast library was incubated with 125 nM of tetrameric SARS-CoV-1 and 1 µl of anti-c-myc FITC (Miltenyi) for 1 hour. Samples were then washed 2x with PBSM and then resuspended in 50 µl PBSM. These libraries were then sorted on an FACSAria IIu using the Stanford FACS Facility (Stanford CA). The samples were gated such that all antigen positive cells were collected (gates set such that ∼0% anti-cmyc FITC alone controls fell within the gate). Two populations were sorted, a hi-gate, consisting of the highest intensity binders (3.8% of all cells), and a low-gate, consisting of all other antigen positive cell (3.7% of all cells). Cells were sorted directly into tubes containing 4mL of SD-CAA media. These sorted libraries were grown for 1 day at 30°C shaking in SD-CAA media and then 300 µl of the cultures were miniprepped (Zymo Research) following the manufacturer’s protocol. Miniprepped DNA was transformed into STELLAR Competent Cells (Clontech) and plated on carbenicillin LB agar plates (as per pPNL6’s resistance marker). E. coli cells that grow should, theoretically, contain only a single sequence from each of the yeast that were sorted above. 10 E. coli colonies from the hi-gate and 20 E. coli colonies from the low-gate sort were sent for sequencing (Sequetech, Mountain View CA). The sequences were then analyzed by sequence alignment using SnapGene software.

### Constructs

#### scFv-ACE2 fusion proteins

scFvs identified as cross-reacting with SARS-CoV-1 and falling into a unique epitope (CV10, CV27, COVA2-14, COV2-2449, COV2-2143) sort were cloned into the pTwist CMV BetaGlobin vector such that they contained a linker (GGSGSHHHHHHASTGGGSGGPSGQAGAAASEENLYFQGSLFVSNHAYGGSGGEARV) followed by the ectodomain of human ACE2.

#### Light chain (LC) and LC-ACE2 fusion proteins

Antibody sequences were cloned into the CMV/R plasmid backbone for expression under a CMV promoter. The antibodies variable LC were cloned between the CMV promoter and the bGH poly(A) signal sequence of the CMV/R plasmid to facilitate improved protein expression. The variable region was cloned into the human IgG1 backbone with a kappa LC. This vector also contained the HVM06_Mouse (P01750) Ig heavy chain V region 102 signal peptide to allow for protein secretion and purification from the supernatant. The light chains from the scFvs from the above-described SARS-CoV-1 sort were cloned into the CMV/R vector in frame with the kappa LC. For COV2-2449 the LC was additionally cloned such that there was a C terminal linker (GGSGSHHHHHHASTGGGSGGPSGQAGAAASEENLYFQGSLFVSNHAYGGSGGEARV)) followed by the ectodomain of human ACE2.

#### Heavy Chain (HC) IgG plasmids

Antibody sequences were cloned into the CMV/R plasmid backbone for expression under a CMV promoter. The antibodies variable HC were cloned between the CMV promoter and the bGH poly(A) signal sequence of the CMV/R plasmid to facilitate improved protein expression. The variable region was cloned into the human IgG1 backbone. This vector also contained the HVM06_Mouse (P01750) Ig heavy chain V region 102 signal peptide to allow for protein secretion and purification from the supernatant. The heavy chains from the scFvs from the above-described SARS-CoV-1 sort were cloned into the CMV/R vector in frame with HC constant regions.

#### hCoV spike proteins constructs

Spike proteins from six hCoVs were cloned into a pADD2 vector between the rBeta-globin intron and β-globin poly(A). A total of 48 constructs were cloned and tested containing either a C-terminal truncation or not, a foldon or GCN4 trimerization domain, and containing an Avi tag or not. Each set of eight proteins was produced for the 6 hCoV spike proteins from SARS-CoV-2, SARS-CoV-1, MERS, 229E, NL63, and OC43. Depictions of the constructs and linkers produced are shown in supplementary Fig. 2.

#### Lentivirus plasmids

Plasmids encoding the full-length spike proteins with native signal peptides were cloned into the background of the HDM-SARS2-Spike-delta21 plasmid (Addgene Plasmid #155130). This construct contains a 21 amino acid c-terminal deletion to promote viral expression. The SARS-CoV-1 spike was used with an 18 amino acid C-terminal deletion. The other viral plasmids that were used were previously described (doi: 10.3390/v12050513). They are: pHAGE-Luc2- IRS-ZsGreen (NR-52516), HDM-Hgpm2 (NR-52517), pRC-CMV-Rev1b (NR-52519), and HDM- tat1b (NR-52518).

#### DNA Preps

The 48 spike protein constructs from the hCoVs were mirA prep using ThermoFisher GeneJET plasmid miniprep kit. 8 mL of *E. Coli* containing the constructs were harvested by centrifugation and 200 µL of freshly made resuspension buffer was added to each clone. 200 µL was then added of lysis buffer followed by inversion and then 300 µL of neutralization buffer was added. Lysed *E. Coli* was then centrifuged at >18,000g for 10 min. The supernatant was transferred to a tube containing 580 µL of 100% EtOH. The EtOH solution was then added to a geneJET plasmid miniprep column and the regular wash steps and elution steps were followed. MirA preps resulted in significantly more plasmid production and allowed for small scale transfection of the 48 clones tested. For the FL-GCN4-Avi-His expression tests and protein production, all samples were maxi prepped from 200 mL of *E. Coli* using NuceloBond Xtra Maxi Kit per the manufacturers recommendations (Macherey-Nagel). All scFv-ACE2, CrossMAb, antibody, hFc-ACE2, and Lentiviral plasmids were maxi prepped in the same fashion.

#### Protein Production

Protein Expression. All proteins were expressed in Expi293F cells. Expi293F cells were cultured in media containing 66% Freestyle/33% Expi media (ThermoFisher) and grown in TriForest polycarbonate shaking flasks at 37°C in 8% CO2. The day before transfection cells were spun down and resuspended to a density of 3x10^6^ cells/mL in fresh media. The following day cells were diluted and transfected at a density of approximately 3-4x10^6^ cells/mL. Transfection mixtures were made by adding the following components: mirA-prepped or maxi-prepped DNA, culture media, and FectoPro (Polyplus) would be added to cells to a ratio of .5-.8µg:100µL:1.3µL:900µL. For example, for a 100mL transfection, 50-80µg of DNA would be added to 10mL of culture media and then 130uL of FectoPro would be added to this. Following mixing and a 10min incubation, the resultant transfection cocktail would be added to 90mL of cells. The cells were harvested 3-5 days post-transfection by spinning the cultures at >7,000 x g for 15 minutes. Supernatants were filtered using a 0.22 µm filter. To determine hCoV protein expression, spun-down Expi293F supernatant was used without further purification. For proteins containing a biotinylation tag (Avi- Tag) Expi293F cells containing a stable BirA enzyme insertion were used, resulting in spontaneous biotinylation during protein expression.

#### Protein purification - Fc Tag containing proteins

All proteins containing an Fc tag (for example, IgGs, CrossMAb-Ace2 fusions, hFc-ACE2) were purified using a 5mL MAb Select Sure PRISM™ column on the AKTA pure FPLC. (Cytiva). Filtered cell supernatants were diluted with 1/10th volume 10x Phosphate Buffered Saline (PBS). The AKTA system was equilibrated with, A1 – 1xPBS, A2 – 100 mM Glycine pH 2.8, B1 – 0.5M NaOH, Buffer line – 1xPBS, Sample lines – H2O. The protocol washes the column with A1, followed by loading of the sample in Sample line 1 until air is detected in the air sensor of the sample pumps, followed by 5 column volume washes with A1, elution of the sample by flowing of 20mL of A2 (directly into a 50mL conical containing 2mL of 1M Tris pH 8.0) followed by 5 column volumes A1, B1, A1. The resultant Fc-containing samples were concentrated using 50 or 100 kDa cutoff centrifugal concentrators. Proteins were buffer exchanged using a PD-10 column (SEPHADEX) which had been preequilibrated into 20 mM HEPES, 150 mM NaCl. IgGs used for competition, binding, and neutralization experiments were not further purified. CrossMAb-ACE2 fusions were then further purified using the S6 column on the AKTA.

#### Protein purification – His-tagged proteins

All proteins not containing an Fc tag (for example, scFvs and scFv fusions, and FL Spike trimers from hCoVs polypeptide antigens) were purified using HisPur™ Ni-NTA resin (ThermoFisher). Cell supernatants were diluted with 1/3rd volume wash buffer (20 mM imidazole, 20 mM HEPES pH 7.4, 150 mM NaCl) and the Ni-NTA resin was added to diluted cell supernatants. For all mixtures not containing hCoV spike protein, the samples were then incubated at 4°C while stirring overnight. hCoV spike proteins were incubated at room temperature. Resin/supernatant mixtures were added to chromatography columns for gravity flow purification. The resin in the column was washed with wash buffer (20 mM imidazole, 20 mM HEPES pH 7.4, 150 mM NaCl) and the proteins were eluted with 250 mM imidazole, 20 mM HEPES pH 7.4, 105mM NaCl. Column elutions were concentrated using centrifugal concentrators (50 kDa cutoff for scFv-ACE2-fusions, and 100 kDa cutoff for trimer constructs), followed by size-exclusion chromatography on a AKTA Pure system (Cytiva). AKTA pure FPLC with a Superdex 6 Increase gel filtration column (S6) was used for purification. 1mL of sample was injected using a 2mL loop and run over the S6 which had been preequilibrated in degassed 20 mM HEPES, 150 mM NaCl prior to use. Biotinylated antigens were not purified using the AKTA pure.

#### TEV Digestion

TEV digestion of scFv-ACE2 fusions. 2µL of TEV protease (New England BioLabs) was added per 200 µL of scFv-ACE2 fusions at ∼4µM in 20 mM HEPES, 150 mM NaCl. The reaction was left to incubate overnight at 30°C. Extent of cleavage was determined by and SDS-PAGE analysis on 4-20% Mini-PROTEAN® TGX^™^ protein gels stained with GelCode^™^ Blue Stain Reagent (ThermoFisher). TEV digestion of CrossMAb-ACE2 fusions. 3µL of TEV protease (New England BioLabs) was added per 200µL of CrossMAb-ACE2 fusions at ∼2µM in 20 mM HEPES, 150 mM NaCl. The reaction was left to incubate overnight at 30°C. Extent of cleavage was determined by and SDS-PAGE analysis on 4-20% Mini-PROTEAN® TGX^™^ protein gels stained with GelCode^™^ Blue Stain Reagent (ThermoFisher).

#### FAb Production from IgGs

1/10 volume of 1M Tris, pH 8 was added to IgGs at ∼2 mg/mL in PBS. 2 µL of a 1 mg/mL stock of Lys-C (stock stored at -70C) was added for each mg of human IgG1 and digested for 1 hour at 37 °C with moderate rotation. Digested FAbs were purified by SP/AKTA using 50 mM NaOAc, pH5 with gradient NaCl elution (using 50 mM NaOAc + 1M NaCl, pH5). FAb fractions were pooled and dialyze against 1x PBS and concentrated using 30kDa concentrators. Purified FAbs were stored at -80 °C.

#### Biolayer Interferometry Binding

Biolayer interferometry (Octet) Binding Experiments – hCoV expression testing. All reactions were run on an Octet Red 96 and samples were run in PBS with 0.1% BSA and 0.05% Tween 20 (octet buffer). hCoVs supernatants were assessed for binding using anti-Penta His (His1K) tips were used. These tips are designed to bind specifically to a penta-His tag on proteins. For this experiment, tips were baselined in a blank well and then associated in the wells containing 50 µL of hCoV expression media and 150 µL octet buffer. Response values (I.E. peak reached after 5 mins of association) was determined using the Octet data analysis software. Final data analysis was done in Prism.

Biolayer interferometry (Octet) Binding Experiments IgG binding. All reactions were run on an Octet Red 96 and samples were run in PBS with 0.1% BSA and 0.05% Tween 20 (octet buffer). IgGs produced from the scFvs from the above sort were assessed for binding using streptavidin (SA) biosensors (Sartorius/ForteBio) loaded to a threshold of 0.8nm of SARS-CoV- 2, SARS-CoV-1, MERS, and OC43 biotinylated spike proteins. Tips were then washed and baselined in wells containing only octet buffer. Samples were then associated in wells containing 100nM IgG. A control well which loaded antigen but associated in a well containing only 200 µL octet buffer was used as a baseline subtraction for data analysis.

Biolayer interferometry (Octet) Binding Experiments IgG competition. All reactions were run on an Octet Red 96 and samples were run in PBS with 0.1% BSA and 0.05% Tween 20 (octet buffer). IgGs produced from the scFvs from the above sort were assessed for their competition of binding with one another using anti-Penta HIS (His1K) biosensors (Sartorius/ForteBio). His1K tips were pre-quenched with buffer containing 10nM biotin. Tips were then loaded with 100nM protein for 2 mins (SARS-CoV-2 spike) or 4 mins (SARS-CoV-1 spike). These tips were then associated with one of seven antibodies (either CV27, COV2-2147, CV10, COVA2-14, COVA2-18, COV2- 2449, COV2-2143) at 100nM for 5mins to reach saturation. Tips were then baselined and associated with either 1 of the 7 antibodies. For this step all 8 tips went into the same antibody at 100nM. Response values (I.E. peak reached after 2 mins of association) was determined using the Octet data analysis software. Values were normalized to the tip loaded with either SARS-CoV- 2 or SARS-CoV-1 spike but without a competing antibody. These values were set as a value of 1 for each antibody. This is simply the antibody binding to the protein. Additionally, the antibody competing with itself was set to a value of zero. Final data analysis was done in Prism.

Biolayer interferometry (Octet) Binding Experiments - scFv-ACE2-Fusion. All reactions were run on an Octet Red 96 and samples were run in PBS with 0.1% BSA and 0.05% Tween 20. Streptavidin (SA) biosensors (Sartorius/ForteBio) were loaded for 2mins with 100 nM biotinylated antigens (SARS-CoV-2 or SARS-CoV-1 spike proteins). Samples were then washed and baselined in wells containing octet buffer. Association occurred in samples containing ACE2- fusion proteins either without or with TEV protease (NEB) treatment. scFv-ACE2 fusions were tested at 200nM. Association was conducted for 2 min and dissociation was conducted for 1 min.

Biolayer interferometry (Octet) Binding Experiments - scFv-ACE2-fusion and CrossMAb-ACE2-fusion competition with hFc-ACE2. All reactions were run on an Octet Red 96 and samples were run in PBS with 0.1% BSA and 0.05% Tween 20 (octet buffer). Streptavidin (SA) biosensors (Sartorius/ForteBio) (scFvs) or HIS 1K biosensors (Sartorius/ForteBio) (CrossMAb) were loaded for 2mins with 100 nM biotinylated antigens (SARS-CoV-2 or SARS-CoV-1 spike – scFvs) or 4 mins with 200 nM his-tagged antigens (CrossMAb). Samples were then washed and baselined in wells containing octet buffer. scFv-ACE2-fusions or CrossMAbs were then associated for 5mins. Samples were baselined and then associated with either hFc-ACE2 for 2mins (scFv) or 40 seconds (CrossMAb). Response values was determined using the Octet data analysis software. Samples which loaded SARS-CoV-2 or SARS-CoV-1 but did not associate with any hFc-ACE2 were used as a baseline subtraction. Values were normalized to the binding of hFc-ACE2 without a competitor.

#### Lentivirus Production

SARS-CoV-2, VOCs, and SARS-CoV-1 spike pseudotyped lentiviral particles were produced. Viral transfections were done in HEK293T cells using calcium phosphate transfection reagent. Six million cells were seeded in D10 media (DMEM + additives: 10% FBS, L-glutamate, penicillin, streptomycin, and 10 mM HEPES) in 10 cm plates one day prior to transfection. A five-plasmid system (plasmids described above) was used for viral production, as described in Crawford et al., 2020. The Spike vector contained the 21 amino acid truncated form of the SARS-CoV-2 Spike sequence from the Wuhan-Hu-1 strain of SARS-CoV-2 or VOCs, or 18 amino acid truncation for SARS-CoV-1. VOCs were based off WT – Sequence ID: BCN86353.1, Alpha – Sequence ID: QXN08428.1, Beta – Sequence ID: QUT64557.1, Gamma – Sequence ID: QTN71704.1, Delta – sequence ID: QWS06686.1 which also has V70F and A222V mutations, and Omicron – sequence ID: UFO69279.1. The plasmids were added to D10 medium in the following ratios: 10 µg pHAGE-Luc2-IRS-ZsGreen, 3.4 µg FL Spike, 2.2 µg HDM-Hgpm2, 2.2 µg HDM-Tat1b, 2.2 µg pRC-CMV-Rev1b in a final volume of 1000 µL. To form transfection complexes, 30 µL BioT (BioLand) was added. Transfection reactions were incubated for 10 min at RT, and then 9 mL of medium was added slowly. The resultant 10mL was added to plated HEK cells from which the medium had been removed. Culture medium was removed 24 hours post-transfection and replaced with fresh D10 medium. Viral supernatants were harvested 72 hours post-transfection by spinning at 300 x g for 5 min followed by filtering through a 0.45 µm filter. Viral stocks were aliquoted and stored at -80°C until further use.

#### Neutralization

The target cells used for infection in viral neutralization assays were from a HeLa cell line stably overexpressing the SARS-CoV-2 receptor, ACE2, as well as the protease known to process SARS-CoV-2, TMPRSS2. Production of this cell line is described in detail in Rogers et al., 2020, with the addition of stable TMPRSS2 incorporation. ACE2/TMPRSS2/HeLa cells were plated one day prior to infection at 5,000 cells per well. 96 well white walled, white bottom plates were used for the assay (Thermo Fisher Scientific). On the day of the assay, purified CrossMAb- or scFv-ACE2 fusions in HEPES (20 mM), NaCl (150 mM), which either had or had not been treated with TEV protease (as above), were sterile filtered using a .22 µm filter. Dilutions of this filtered stock were made into sterile 1xDPBS (Thermo Fisher Scientific) which was 5% by volume D10 medium. Each dilution well contained 30 µL of CrossMAb- or scFv-ACE2 fusions. Samples were run in technical duplicate in each experiment. Virus only wells and cell only wells other wells contained only 30 µL 1xDPBS.

A virus mixture was made containing the virus of interest (for example SARS-CoV-2), and D10 media (DMEM + additives: 10% FBS, L-glutamate, penicillin, streptomycin, and 10 mM HEPES). Virus dilutions into media were selected such that a suitable signal would be obtained in the virus only wells. A suitable signal was selected such that the virus only wells would achieve a luminescence of at least >10,000 RLU. 90 µL of this virus mixture was added to each of the inhibitor dilutions to make a final volume of 120 µL in each well. Virus only wells were made which contained 30 µL 1xDPBS and 90 µL virus mixture. Cells only wells were made which contained 30 µL 1xDPBS and 90 µL D10 media.

The inhibitor/virus mixture was left to incubate for 1 hour at 37°C. Following incubation, the medium was removed from the cells on the plates made 1 day prior. This was replaced with 100 µL of inhibitor/virus dilutions and incubated at 37°C for approximately 24 hours. At 24 hours post infection the media was exchanged for fresh media in all samples containing a TEV cleavable linker with our without cleavage, media was not exchanged on samples which did not have a TEV cleavable linker (for example WT IgGs). Infectivity readout was performed by measuring luciferase levels. 48 hours post infection 50 µL of medium was removed from all cells were lysed by the addition of 50 µL BriteLite™ assay readout solution (Perkin Elmer) into each well, alternatively, all the media was removed and a 1:1 dilution of BriteLite™ was used. Luminescence values were measured using a BioTek Synergy™ HT Microplate Reader (BioTek) plate reader. Each plate was normalized by averaging cells only (0% infectivity) and virus only (100% infectivity) wells. Cells only and virus only wells were averaged. Normalized values were fit with a four parameter non-linear regression inhibitor curve in Prism to obtain IC50 values. The average NT_50_ of two independent experiments are shown.

### ELISA

IgG ELISAs against hCoV strains were performed. Streptavidin solution (5 µg/mL) was plated in 50µL in each well on a MaxiSorp (Thermo Fisher Scientific) microtiter plate in 50 mM sodium bicarbonate pH 8.75. This was left to incubate for 1 hour at room temperature. These were washed 3x with 300 µL of ddH2O using an ELx 405 Bio-Tex plate washer and blocked with 150uL Chonblock (Chondrex) for at least 1 hour at room temperature. Biotinylated hCoV spike proteins were added to each well at a concentration of 1µg/mL and left to incubate overnight at 4°C. Plates were washed 3x with 300 µL of 1xPBST and serial dilution of monoclonal antibodies (described above) were added, starting at 1µM and undergoing 10-fold serial dilutions. These were left to incubate for 1 hour at room temperature and then washed 3x with PBST. Goat anti-human HRP (Abcam ab7153) was added at a 1:5,000 dilution in PBST. This was left to incubate at room temperature for 1 hour and then washed 6x with PBST. Finally, the plate was developed using 50 µL of 1-StepTM Turbo-TMB-ELISA Substrate Solution (ThermoFisher) per well and the plates were quenched with 50 µL of 2M H_2_SO_4_ to each well. Plates were read at 450 nm and normalized for path length using a BioTek Synergy™ HT Microplate Reader.

#### Dot blot analysis of hCoV expression

Expi293F culture supernatants from hCoV spike antigen expressions using MirA preps as above were harvested 2 days post-transfection via centrifugation at 7000g for 15 min. Supernatants were spotted on a nitrocellulose membrane The blot was dried for 15 min in a fume hood. Following drying, 10 mL of 1x PBST + 5% blotting grade blocker (Bio-Rad) were added for 10 min. Two microliters of mouse anti-hexa His antibody (BioLegend) were added to the 10 mL sample and incubated for 1 h at room temperature. Blots were washed 16 times with 9 mL of PBST. Ten milliliters of 1x PBST + 5% blotting grade blocker with 2 μL anti-mouse IgG1 (Abcam) were added and incubated for 1 h at room temperature. Blots were washed 16 times with 9 mL of PBST, developed using Pierce ECL Western blotting substrate, and imaged using a GE Amersham imager 600.

## Supplementary Information

### Sequences

**<u>Signal Peptide</u>**

<u>Hexa-His tag</u>

**TEV Site**

<u>ACE2</u>

#### CV10-ACE2 Fusion scFv

**<u>MGWSCIILFLVATATGVHS</u>**QVQLQESGPGLVKPSETLSLTCNVSGGSISSYYWSWIRQP PGKGLEWIGYIYYSGSTNYNPSLKSRVTISVDTSKNQFSLKLSSVTAADTAVYYCARGFDYWGQGTLVTVSSASGGGGSGGGGSGGGGSEIVLTQSPGTLSLSPGERATLSCRASQSVSSIYLAWYQQKPGQAPRLLIYGASSRATGIPDRFSGSGSGTDFTLTISRLEPEDFAVYYCQQ YAGSPWTFGQGTKVEIKGGSGS*<u>HHHHHH</u>*ASTGGGSGGPSGQAGAAASE**ENLYFQG**SLF VSNHAYGGSGGEARV<u>STIEEQAKTFLDKFNHEAEDLFYQSSLASWNYNTNITEENVQN MNNAGDKWSAFLKEQSTLAQMYPLQEIQNLTVKLQLQALQQNGSSVLSEDKSKRLNTILNTMSTIYSTGKVCNPDNPQECLLLEPGLNEIMANSLDYNERLWAWESWRSEVGKQLRPLYEEYVVLKNEMARANHYEDYGDYWRGDYEVNGVDGYDYSRGQLIEDVEHTFEEIKPLYEHLHAYVRAKLMNAYPSYISPIGCLPAHLLGDMWGRFWTNLYSLTVPFGQKPNID VTDAMVDQAWDAQRIFKEAEKFFVSVGLPNMTQGFWENSMLTDPGNVQKAVCHPTAWDLGKGDFRILMCTKVTMDDFLTAHHEMGHIQYDMAYAAQPFLLRNGANEGFHEAVGEIMSLSAATPKHLKSIGLLSPDFQEDNETEINFLLKQALTIVGTLPFTYMLEKWRWMVFKGEIPKDQWMKKWWEMKREIVGVVEPVPHDETYCDPASLFHVSNDYSFIRYYTRTLYQFQFQEALCQAAKHEGPLHKCDISNSTEAGQKLFNMLRLGKSEPWTLALENVVGAKNMNVRPLLNYFEPLFTWLKDQNKNSFVGWSTDWSPYAD</u>

#### CV27-ACE2 Fusion scFv

**<u>MGWSCIILFLVATATGVHS</u>**QVQLVESGGGVVQPGRSLRLSCAASGFTFSSYAMHWVRQAPGKGLEWVALISYDGSNKYYADSVKGRFTISRDNSKNTLYLQMNSLRAEDTAVYYC ARSFGGSYYYGMDVWGQGTTVTASGGGGSGGGGSGGGGSQSALTQPASVSGSPGQSITISCTGTSSDVGGYNYVSWYQQHPGKAPKLMIYDVSNRPSGVSNRFSGSKSGNTASLTIS GLQAEDEADYYCSSYTSSSTPYVFGTGTKVGGSGS*<u>HHHHHH</u>*ASTGGGSGGPSGQAGAAASE**ENLYFQG**SLFVSNHAYGGSGGEARV<u>STIEEQAKTFLDKFNHEAEDLFYQSSLASWNYNTNITEENVQNMNNAGDKWSAFLKEQSTLAQMYPLQEIQNLTVKLQLQALQQNGSSVLSEDKSKRLNTILNTMSTIYSTGKVCNPDNPQECLLLEPGLNEIMANSLDYNERLWAWESWRSEVGKQLRPLYEEYVVLKNEMARANHYEDYGDYWRGDYEVNGVDGYDYSRGQLIEDVEHTFEEIKPLYEHLHAYVRAKLMNAYPSYISPIGCLPAHLLGDMWGRFWTNLYSLTVPFGQKPNIDVTDAMVDQAWDAQRIFKEAEKFFVSVGLPNMTQGFWENSMLTDPGNVQKAVCHPTAWDLGKGDFRILMCTKVTMDDFLTAHHEMGHIQYDMAYAAQPFLLRNGANEGFHEAVGEIMSLSAATPKHLKSIGLLSPDFQEDNETEINFLLKQALTIVGTLPFTYMLEKWRWMVFKGEIPKDQWMKKWWEMKREIVGVVEPVPHDETYCDPASLFHVSNDYSFIRYYTRTLYQFQFQEALCQAAKHEGPLHKCDISNSTEAGQKLFNMLRLGKSEPWTLALENVVGAKNMNVRPLLNYFEPLFTWLKDQNKNSFVGWSTDWSPYAD</u>

#### COVA2-14-ACE2 Fusion scFv

**<u>MGWSCIILFLVATATGVHS</u>**QVQLVQSGAEVKKPGSSVKVSCKASGGTFSSYAIIWVRQAPGQGLEWMGGIIPIFGTANYAQKFQGRVTITTDESTSTAYMELSSLRSEDTAVYYCAR VRYYDSSGYYEDYWGQGTLVTVSSASGGGGSGGGGSGGGGSEIVLTQSPATLSLSPGERATLSCRASQSVSSYLAWYQQEPGQAPRLLIYDASNRATGIPARFSGSGSGTDFTLTISSLE PEDFAVYYCQQRSNWPPMYTFGQGTKVEIKGGSGS*<u>HHHHHH</u>*ASTGGGSGGPSGQAGAAASE**ENLYFQG**SLFVSNHAYGGSGGEARV<u>STIEEQAKTFLDKFNHEAEDLFYQSSLASWNYNTNITEENVQNMNNAGDKWSAFLKEQSTLAQMYPLQEIQNLTVKLQLQALQQNGSSVLSEDKSKRLNTILNTMSTIYSTGKVCNPDNPQECLLLEPGLNEIMANSLDYNERLWAWESWRSEVGKQLRPLYEEYVVLKNEMARANHYEDYGDYWRGDYEVNGVDGYDYSRGQLIEDVEHTFEEIKPLYEHLHAYVRAKLMNAYPSYISPIGCLPAHLLGDMWGRFWTNLYSLTVPFGQKPNIDVTDAMVDQAWDAQRIFKEAEKFFVSVGLPNMTQGFWENSMLTDPGNVQKAVCHPTAWDLGKGDFRILMCTKVTMDDFLTAHHEMGHIQYDMAYAAQPFLLRNGANEGFHEAVGEIMSLSAATPKHLKSIGLLSPDFQEDNETEINFLLKQALTIVGTLPFTYMLEKWRWMVFKGEIPKDQWMKKWWEMKREIVGVVEPVPHDETYCDPASLFHVSNDYSFIRYYTRTLYQFQFQEALCQAAKHEGPLHKCDISNSTEAGQKLFNMLRLGKSEPWTLALENVVGAKNMNVRPLLNYFEPLFTWLKDQNKNSFVGWSTDWSPYAD</u>

#### COV2-2449-ACE2 Fusion scFv

**<u>MGWSCIILFLVATATGVHS</u>**QVQLVESGGGVVQPGRSLRLSCATSGFTFSSFALHWVRQAPGKGLEWVTVISDDGNNKYYVDSVKGRFTISRDNSKNTLFLQMNSLRVEDTAIYYCA RASYNSNWSIGEYFRDWGQGTLVTVSSASGGGGSGGGGSGGGGSDIVMTQSPDSLAVSLGERATINCKSSQSLLYTSNNKNYLAWYQQKPGQPPKLLIYWASTRESGVPDRFSGSGS GTDFTLTISSLQAEDVAVYYCQQYYSPPWTFGQGTKVEIKGGSGS*<u>HHHHHH</u>*ASTGGGSGGPSGQAGAAASE**ENLYFQG**SLFVSNHAYGGSGGEARV<u>STIEEQAKTFLDKFNHEAEDLFYQSSLASWNYNTNITEENVQNMNNAGDKWSAFLKEQSTLAQMYPLQEIQNLTVKLQLQALQQNGSSVLSEDKSKRLNTILNTMSTIYSTGKVCNPDNPQECLLLEPGLNEIMANSLDYNERLWAWESWRSEVGKQLRPLYEEYVVLKNEMARANHYEDYGDYWRGDYEVNGVDGYDYSRGQLIEDVEHTFEEIKPLYEHLHAYVRAKLMNAYPSYISPIGCLPAHLLGDMWGRFWTNLYSLTVPFGQKPNIDVTDAMVDQAWDAQRIFKEAEKFFVSVGLPNMTQGFWENSMLTDPGNVQKAVCHPTAWDLGKGDFRILMCTKVTMDDFLTAHHEMGHIQYDMAYAAQPFLLRNGANEGFHEAVGEIMSLSAATPKHLKSIGLLSPDFQEDNETEINFLLKQALTIVGTLPFTYMLEKWRWMVFKGEIPKDQWMKKWWEMKREIVGVVEPVPHDETYCDPASLFHVSNDYSFIRYYTRTLYQFQFQEALCQAAKHEGPLHKCDISNSTEAGQKLFNMLRLGKSEPWTLALENVVGAKNMNVRPLLNYFEPLFTWLKDQNKNSFVGWSTDWSPYAD</u>

#### COV2-2143-ACE2 Fusion scFv

**<u>MGWSCIILFLVATATGVHS</u>**EVQLVESGGGLVQPGGSLRLSCAASGFTVSSNYMSWVRQAPGKGLEWVSVIYSAGSTYYADSVKGRFSISRDKSKNTLYLQMNSLRAEDTAVYYCA KEGGSGSLRYYYYGMDVWGQGTTVTVSSASGGGGSGGGGSGGGGSQSVVTQPPSASGTPGQRVTISCSGSSSNIGYNIVNWYQQLPGTAPKLLIYSNNQRPSGVPDRFSGSKSGTSAS LSISGLQSEDEADYYCAAWDDSLNGYVFGTGTKVTVLGGSGS*<u>HHHHHH</u>*ASTGGGSGGPSGQAGAAASE**ENLYFQG**SLFVSNHAYGGSGGEARV<u>STIEEQAKTFLDKFNHEAEDLFYQSSLASWNYNTNITEENVQNMNNAGDKWSAFLKEQSTLAQMYPLQEIQNLTVKLQLQALQQNGSSVLSEDKSKRLNTILNTMSTIYSTGKVCNPDNPQECLLLEPGLNEIMANSLDYNERLWAWESWRSEVGKQLRPLYEEYVVLKNEMARANHYEDYGDYWRGDYEVNGVDGYDYSRGQLIEDVEHTFEEIKPLYEHLHAYVRAKLMNAYPSYISPIGCLPAHLLGDMWGRFWTNLYSLTVPFGQKPNIDVTDAMVDQAWDAQRIFKEAEKFFVSVGLPNMTQGFWENSMLTDPGNVQKAVCHPTAWDLGKGDFRILMCTKVTMDDFLTAHHEMGHIQYDMAYAAQPFLLRNGANEGFHEAVGEIMSLSAATPKHLKSIGLLSPDFQEDNETEINFLLKQALTIVGTLPFTYMLEKWRWMVFKGEIPKDQWMKKWWEMKREIVGVVEPVPHDETYCDPASLFHVSNDYSFIRYYTRTLYQFQFQEALCQAAKHEGPLHKCDISNSTEAGQKLFNMLRLGKSEPWTLALENVVGAKNMNVRPLLNYFEPLFTWLKDQNKNSFVGWSTDWSPYAD</u>

**<u>Signal Peptide</u>**

*<u>Hexa-His tag</u>*

**TEV Site**

<u>ACE2</u>

#### COV2-2449-LC-ACE2 Fusion

**<u>MGWSCIILFLVATATGVHS</u>**DIVMTQSPDSLAVSLGERATINCKSSQSLLYTSNNKNYLAWYQQKPGQPPKLLIYWASTRESGVPDRFSGSGSGTDFTLTISSLQAEDVAVYYCQQYYS PPWTFGQGTKVEIKRTVAAPSVFIFPPSDEQLKSGTASVVCLLNNFYPREAKVQWKVDN ALQSGNSQESVTEQDSKDSTYSLSSTLTLSKADYEKHKVYACEVTHQGLSSPVTKSFNR GECGGSGS*<u>HHHHHH</u>*ASTGGGSGGPSGQAGAAASE**ENLYFQG**SLFVSNHAYGGSGGEARV<u>STIEEQAKTFLDKFNHEAEDLFYQSSLASWNYNTNITEENVQNMNNAGDKWSAFLKEQSTLAQMYPLQEIQNLTVKLQLQALQQNGSSVLSEDKSKRLNTILNTMSTIYSTGKVCNPDNPQECLLLEPGLNEIMANSLDYNERLWAWESWRSEVGKQLRPLYEEYVVLKNEMARANHYEDYGDYWRGDYEVNGVDGYDYSRGQLIEDVEHTFEEIKPLYEHLHAYVRAKLMNAYPSYISPIGCLPAHLLGDMWGRFWTNLYSLTVPFGQKPNIDVTDAMVDQAWDAQRIFKEAEKFFVSVGLPNMTQGFWENSMLTDPGNVQKAVCHPTAWDLGKGDFRILMCTKVTMDDFLTAHHEMGHIQYDMAYAAQPFLLRNGANEGFHEAVGEIMSLSAATPKHLKSIGLLSPDFQEDNETEINFLLKQALTIVGTLPFTYMLEKWRWMVFKGEIPKDQWMKKWWEMKREIVGVVEPVPHDETYCDPASLFHVSNDYSFIRYYTRTLYQFQFQEALCQAAKHEGPLHKCDISNSTEAGQKLFNMLRLGKSEPWTLALENVVGAKNMNVRPLLNYFEPLFTWLKDQNKNSFVGWSTDWSPYAD</u>

**<u>Signal Peptide</u>**

***<u>Knob/Hole Mutations</u>***

<u>CrossMAb Swaps</u>

#### COV2-2449-HC-Knob

**<u>MGWSCIILFLVATATGVHS</u>**QVQLVESGGGVVQPGRSLRLSCATSGFTFSSFALHWVRQAPGKGLEWVTVISDDGNNKYYVDSVKGRFTISRDNSKNTLFLQMNSLRVEDTAIYYCA RASYNSNWSIGEYFRDWGQGTLVTVSSASTKGPSVFPLAPSSKSTSGGTAALGCLVKDYFPEPVTVSWNSGALTSGVHTFPAVLQSSGLYSLSSVVTVPSSSLGTQTYICNVNHKPSNTKVDKKVEPKSCDKTHTCPPCPAPELLGGPSVFLFPPKPKDTLMISRTPEVTCVVVDVSHEDPEVKFNWYVDGVEVHNAKTKPREEQYNSTYRVVSVLTVLHQDWLNGKEYKCKVSN KALPAPIEKTISKAKGQPREPQVYTLPPSRDELTKNQVSL***<u>W</u>***CLVKGFYPSDIAVEWESNGQPENNYKTTPPVLDSDGSFFLYSKLTVDKSRWQQGNVFSCSVMHEALHNHYTQKSLSL SPGK

#### CV10-LC-CrossMAb

**<u>MGWSCIILFLVATATGVHS</u>**EIVLTQSPGTLSLSPGERATLSCRASQSVSSIYLAWYQQKPGQAPRLLIYGASSRATGIPDRFSGSGSGTDFTLTISRLEPEDFAVYYCQQYAGSPWTFGQ GTKVEIK<u>SSASTKGPSVFPLAPSSKSTSGGTAALGCLVKDYFPEPVTVSWNSGALTSGVHTFPAVLQSSGLYSLSSVVTVPSSSLGTQTYICNVNHKPSNTKVDKKVEPKSC</u>

#### CV10-HC-CrossMAb-Hole

**<u>MGWSCIILFLVATATGVHS</u>**QVQLQESGPGLVKPSETLSLTCNVSGGSISSYYWSWIRQPPGKGLEWIGYIYYSGSTNYNPSLKSRVTISVDTSKNQFSLKLSSVTAADTAVYYCARGFD YWGQGTLVTVSSAS<u>VAAPSVFIFPPSDEQLKSGTASVVCLLNNFYPREAKVQWKVDNALQSGNSQESVTEQDSKDSTYSLSSTLTLSKADYEKHKVYACEVTHQGLSSPVTKSFNRGEC</u>DKTHTCPPCPAPELLGGPSVFLFPPKPKDTLMISRTPEVTCVVVDVSHEDPEVKFNWYVDGVEVHNAKTKPREEQYNSTYRVVSVLTVLHQDWLNGKEYKCKVSNKALPAPIEKTI SKAKGQPREPQV***<u>C</u>***TLPP***<u>S</u>***RDELTKNQVSL***<u>S</u>***CAVKGFYPSDIAVEWESNGQPENNYKTTPPVLDSDGSFFL***<u>V</u>***SKLTVDKSRWQQGNVFSCSVMHEALHNHYTQKSLSLSPGK

---

**Supplementary Figure 1.**
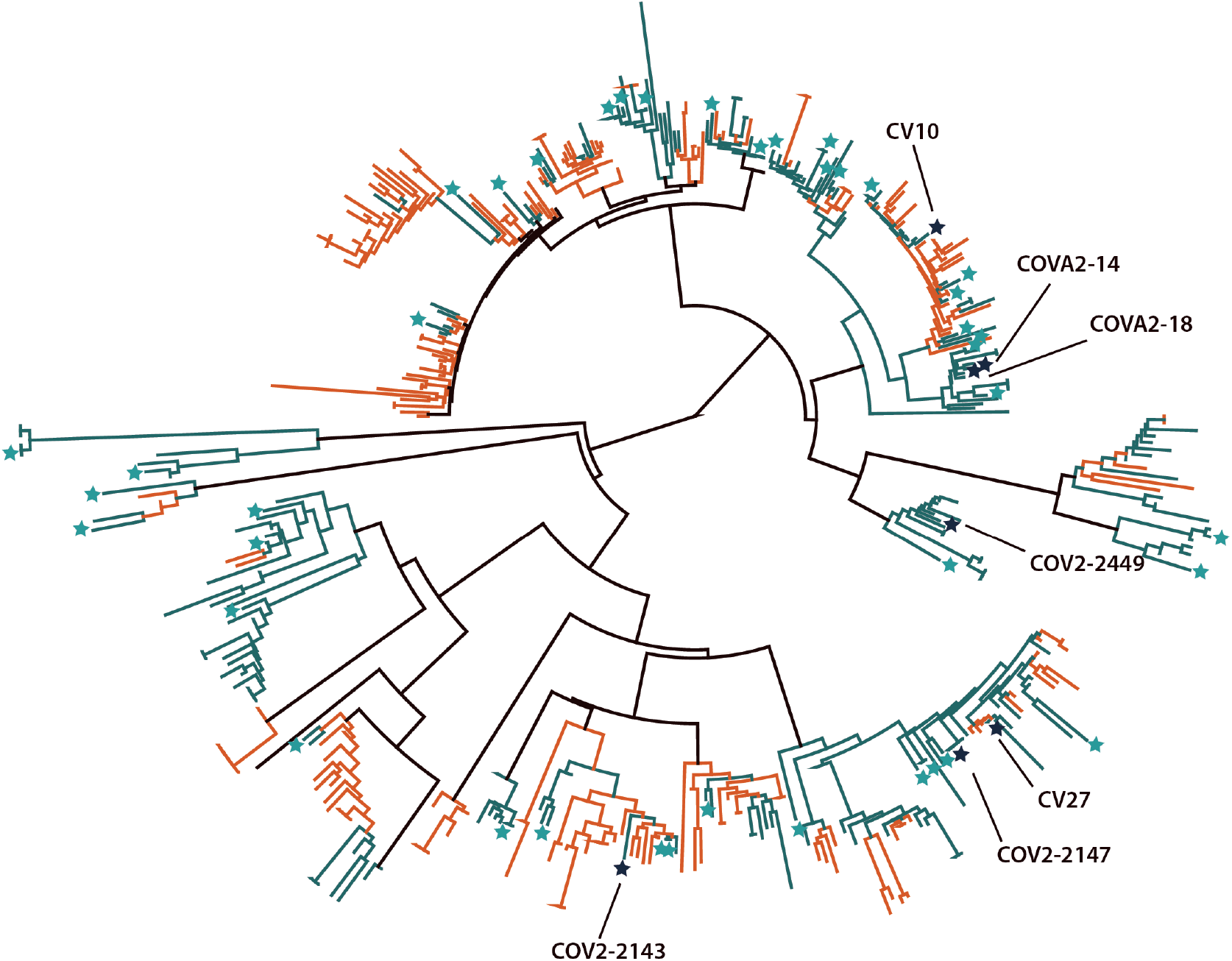
The non-RBD library was selected to prioritize diversity. A phylogenetic tree generated using Geneious Prime of 436 light chain sequences from a curated library of 696 anti-SARS-COV-2 spike antibody sequences. Labels same as Fig. 1. Germline alleles are not shown. Antibodies denoted with names were cross-reactive for SARS-CoV-1.

**Supplementary Figure 2.**
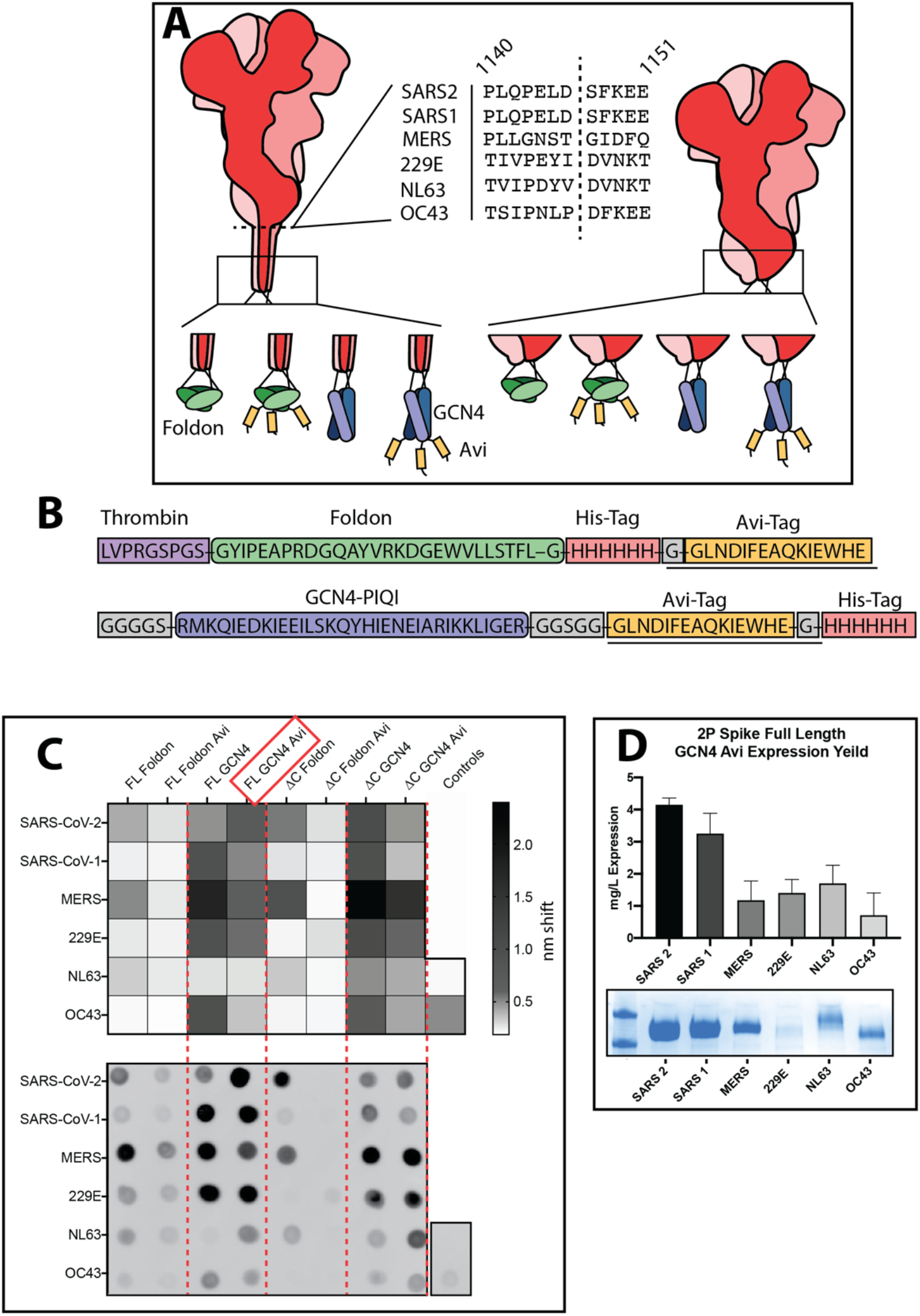
hCoV proteins expression is optimized with a GCN4 tag. (A) A schematic representation of constructs tested for expression. 6 hCoV spike proteins, either FL or truncated, tested with 4 different C-terminal trimerization domains and tags. ΔC truncations shown in the middle, SARS-CoV-2 spike numbering. (B) linkers tested. For those without Avi tags, the underlined portions were removed. (C) (top) BLI binding responses from isolated Expi supernatants binding to His 1K octet tips. Higher response corresponds to higher protein expression. (bottom) An anti-his tag dot blot assay of the supernatants tested in the BLI binding. Dot blot shows good correspondence with the BLI binding. Influenza hemagglutinin used as a positive control, mock transfection as a negative control. (D) Yield determined for full length 2P hCoV constructs containing the GCN4-Avi-His tag from a duplicate experiment of 100 mL transfections.

**Supplementary Figure 3.**
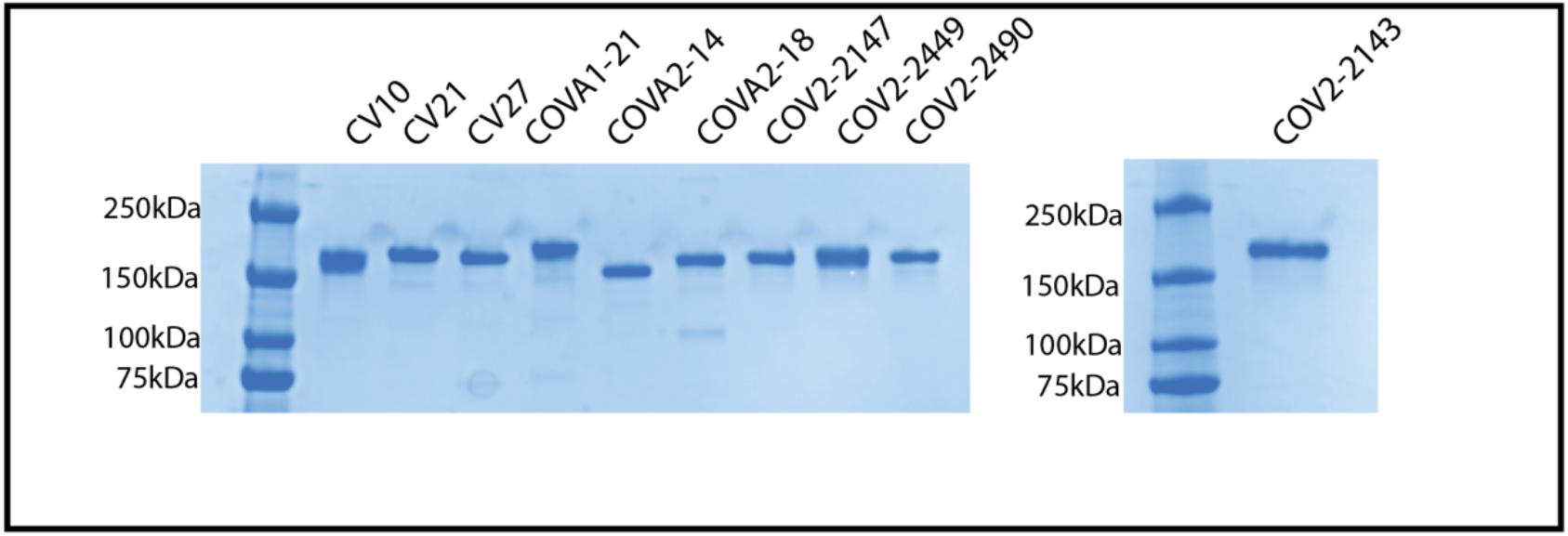
SDS-page analysis of of IgG proteins made from clones identified in the SARS-CoV-1 FACS sorts. MW ladders are in the left-most lanes of the two gels.

**Supplementary Figure 4.**
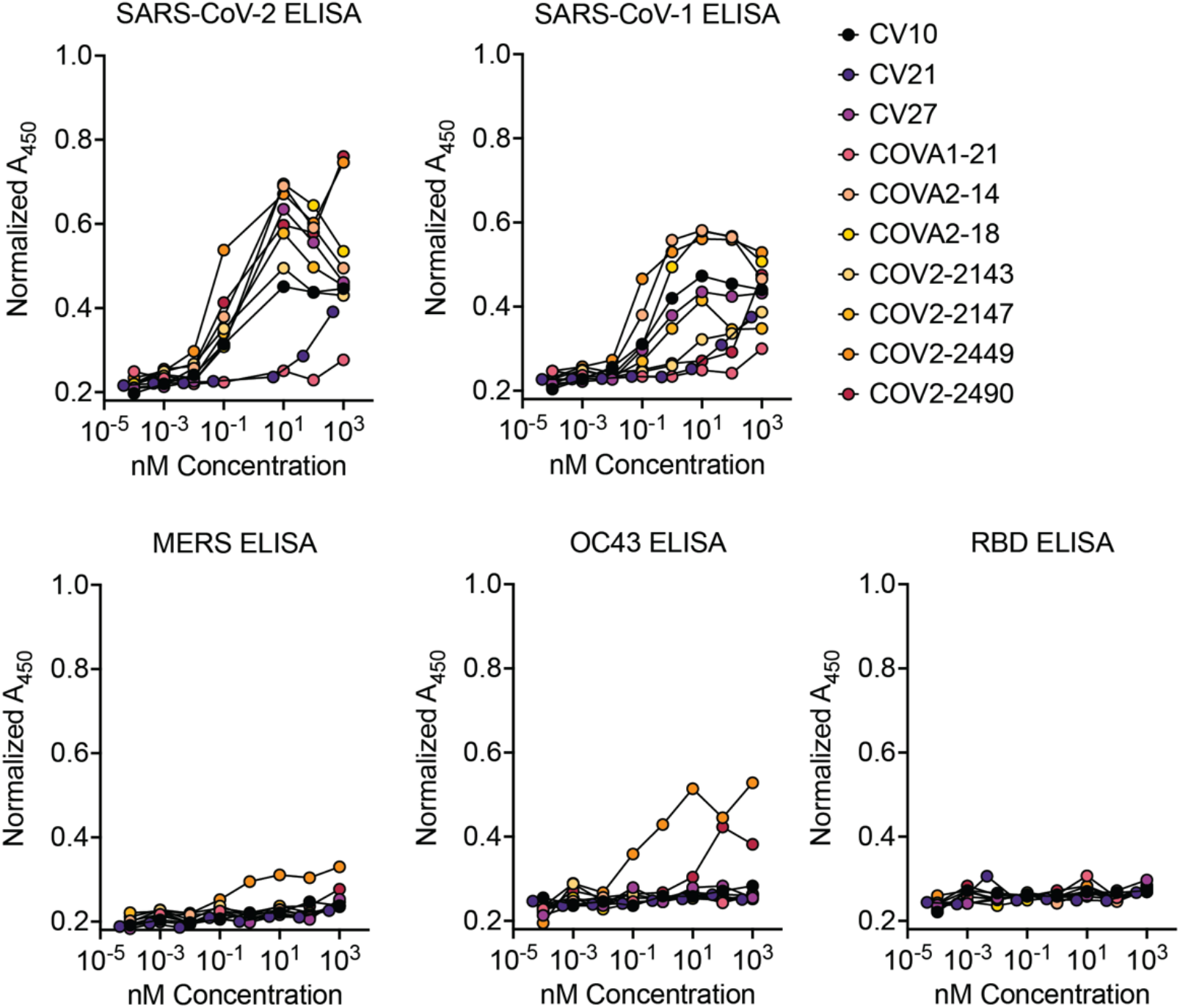
ELISA binding of IgG proteins made from scFv clones identified by FACS sorting for SARS-CoV-1 binding. Biotinylated hCoV antigens were plated and dilutions of IgGs were tested for binding. Normalized A450 calculated by adjusting for pathlength. Except for COV2-2490, CV21, COVA2-18 all IgGs bind to SARS-CoV-2 and SARS-CoV-1.

**Supplementary Figure 5.**
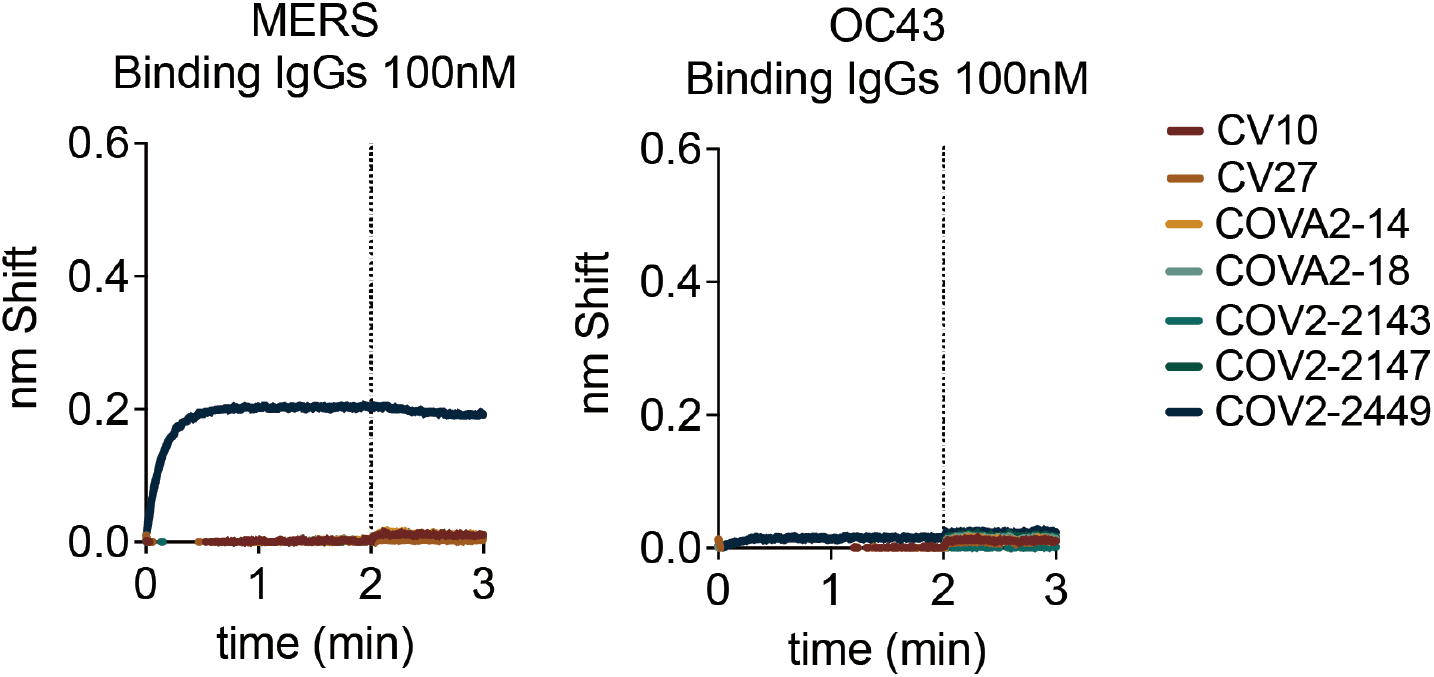
BLI binding for the 7 identified SARS-CoV-1 cross-reactive clones against MERS (left) or OC43 (right) spike proteins. Only COV2-2449 shows any binding affinity for MERS or weakly to OC43 spike proteins.

**Supplementary Figure 6.**
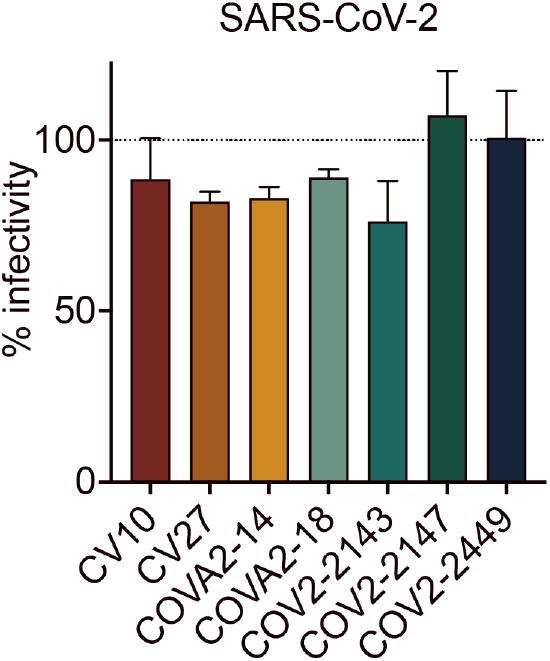
Single dilution (100 nM) neutralization against SARS-CoV-2 Wuhan-Hu-1 for the 7 identified cross-reactive antibodies. All antibodies show no neutralization at a high concentration of 100nM.

**Supplementary Figure 7.**
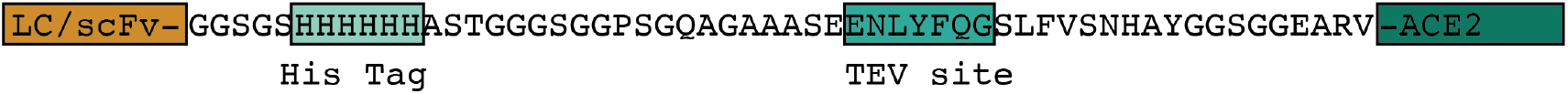
The linker used to tether ACE2 to either the C-terminus of the scFv or C-terminus of the COV2-2449 LC. The linker was designed to contain a Hexa-His tag for purification and a TEV site to facilitate proteolysis.

**Supplementary Figure 8.**
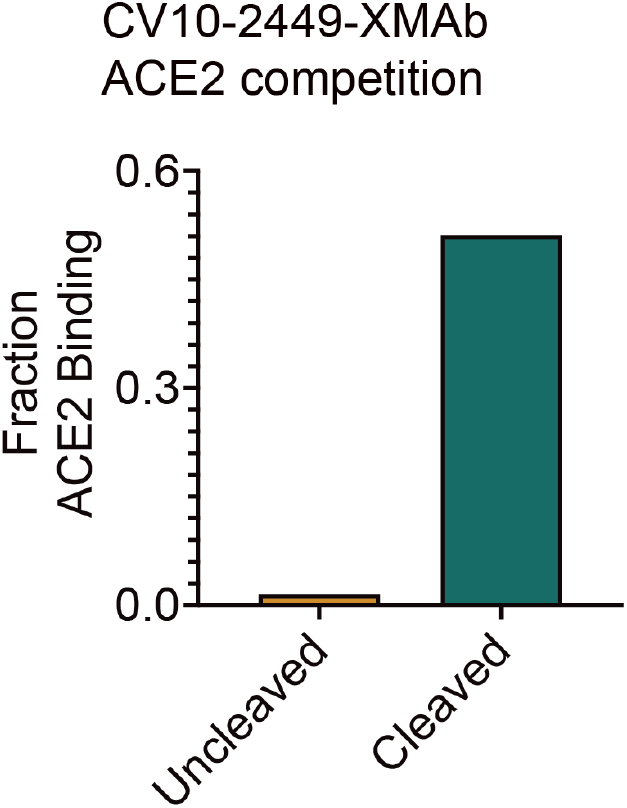
Relative hFc-ACE2 binding to SARS-CoV-2 spike protein which has been pre-associated with 200 nM CV10-2449-XMAb (left) or the TEV cleaved form (right). No competitor was set to a value of 1.0.

